# How infant brains fold: Sulcal deepening is linked to development of sulcal span, thickness, curvature, and microstructure

**DOI:** 10.1101/2025.04.18.649604

**Authors:** Sarah S. Tung, Xiaoqian Yan, Bella Fascendini, Christina Tyagi, Charleny Martinez Reyes, Keithan Ducre, Karla Perez, Ahmad Allen, Juliet Horenziak, Hua Wu, Boris Keil, Vaidehi S. Natu, Kalanit Grill-Spector

## Abstract

Cortical folding begins in utero as sulci emerge and continues postnatally as sulci deepen. However, the timeline and mechanisms underlying postnatal sulcal development remain unknown. Using structural and quantitative magnetic resonance imaging in infants from birth to one year of age, we longitudinally measured macroanatomical and microstructural development in major sulci that emerge in utero between the 16th and 31st gestational weeks. We find that sulci that emerge earlier in utero are deeper at birth and deepen at a slower rate postnatally than later emerging sulci. Sulci also become wider, thicker, and microstructurally denser, while their curvature decreases. Notably, mean sulcal depth is predicted by a weighted sum of sulcal span, thickness, curvature, and tissue microstructure, with differential weights across sulci. Analysis of local depth along the sulcus also reveals that deeper portions of sulci (fundi) have higher curvature and higher microstructural density than the surrounding sulcal walls. These data reveal that postnatal sulcal deepening is nonuniform and depends on the time of emergence in utero, tissue microstructure, and multiple macroanatomical factors. Together, these findings have important ramifications for theories of cortical folding and elucidating neurodevelopmental disorders and delays.

## Introduction

Sulcal folding is a complex process that begins in utero and involves the development of several genetic, macroanatomical, and microstructural features in the brain^1–8^. Early occurring sulcal folds emerge as little valleys by the 14-16^th^ gestational week^1,2,9^ and continue to deepen through the first two years of life^10,11^. One defining morphological measure of sulcal folds that change throughout prenatal and postnatal stages of human life is sulcal depth. Depth is computed as the distance between the cortical surface and the cerebral hull. Importantly, sulcal depth serves as a predictor of functional specialization^12–15^ and irregularities in sulcal depth are biomarkers of neurological disorders like Down syndrome^16^, autism^17^, and depression^18^. Despite the importance of understanding the development of sulcal depth, how sulci deepen during the first year of postnatal life and whether their development is linked with other structural changes, remains unknown. Here, we address this gap in knowledge by using structural and quantitative MRI in infants to examine the development of twelve, early emerging sulci from birth to one year of life.

Cortical development during infancy has garnered much attention over the past decade with the advent of large infant data sets. Macrostructural measures such as gyrification index^19^, sulcal fundi^10,11,20^ (deepest points along a sulcal fold), cortical thickness^21,22^, sulcal depth^23,24^, and mean curvature^25,26^ have been widely used to quantify large-scale cortical folding patterns across the entire brain. While these studies provide valuable insights into global or regional trajectories of cortical development, many analyses have been conducted cross-sectionally and at the group level, limiting our understanding of fine-grained sulcal development over time. Thus, examining the longitudinal development of individual sulci in infants is important for several reasons: First, different sulcal folds emerge at varying gestational stages in utero, raising the question of whether sulci that emerge earlier deepen at different rates than those that emerge later. For example, the calcarine sulcus, which forms around the 16th gestational week, may follow a different developmental trajectory than the occipital-temporal sulcus, which emerges later, around the 28th gestational week^1,2^. Second, sulcal deepening does not occur in isolation but alongside broader cortical expansion, including increases in cortical thickness, surface area, sulcal length, and volume, as well as decreases in cortical curvature^27,28^. Understanding whether these changes are systematically related to sulcal depth is crucial for building a mechanistic model of cortical folding. Third, postnatal cortical maturation involves extensive microstructural changes, including synaptic growth^29,30^, dendritic arborization^31,32^, and myelination^33^, all of which influence tissue density. Advances in quantitative MRI now enable in-vivo measurements of R_1_ (tissue relaxation rate)^34–36^, which is related to tissue density and myelin content, with higher R_1_ values reflecting denser cortical tissue^37–39^. Recent longitudinal measurements of R_1_ in infants reveal regional developmental gradients, with primary visual cortex exhibiting higher R_1_ at birth but slower maturation compared to higher-order visual areas^40^. Despite these insights, it remains unknown how macroanatomical and microstructural changes interact within individual sulci and whether sulcal depth is linked to these concurrent developmental processes.

Although several theories propose that sulcal deepening is related to cortical expansion via increases in surface area and thickness^4,8,41–43^, no study has systematically examined the relation between multiple macro- and microstructural changes within individual sulci during infancy. Here, we examine potential developmental relationships between sulcal depth (SD, **Fig. 1a**) and key structural features (**Fig. 1b**): sulcal span (SP, defined as total sulcal surface area divided by sulcal length; see **Methods** and **Supplementary Fig. 1**), cortical thickness (CT), curvature (CU), and tissue relaxation rate (R_1_). **Sulcal depth vs. Sulcal span (SD-SP)**: One possibility is that cortical expansion in the tangential direction (parallel to the cortical surface) stretches the sulcus laterally, increasing both its span and depth by pushing the sulcal fold deeper into white matter^44^ (**Fig. 1b**, top-left), predicting a positive relationship between SD and SP. Alternatively, sulcal depth may increase without concurrent widening, predicting no relationship between SD and SP (**Fig. 1b**, top-right). **Sulcal depth vs. Cortical thickness (SD-CT):** Radial cortical growth (perpendicular to the cortical surface) could push the sulcus deeper into white matter, leading to simultaneous increases in depth and thickness^8^ (**Fig. 1b**, second row-left), predicting a positive relationship between SD and CT. Alternatively, depth may increase without substantial changes in thickness (**Fig. 1b**, second row-right), suggesting different underlying mechanisms. **Sulcal depth vs Curvature (SD-CU)**: As cortex expands and sulci deepen, their base may widen, decreasing their overall mean curvature and predicting a negative relationship between SD and CU (**Fig. 1b**, third row-left). Alternatively, the sulcus may deepen by simply pushing deeper into white matter while maintaining the mean curvature, predicting no relationship between SD and CU (**Fig. 1b**, third row-right). **Sulcal depth vs Cortical Microstructures (SD-R_1_)**: Increased tissue growth and myelination could exert mechanical, stresses^4^ that deepen the sulcus, predicting a positive relationship between SD and R_1_ relationship (**Fig. 1b**, bottom-left). Conversely, if sulcal depth increases independently of microstructural changes, R_1_ values may remain stable even as sulci deepen (**Fig. 1b**, bottom-right).

**Figure 1.**
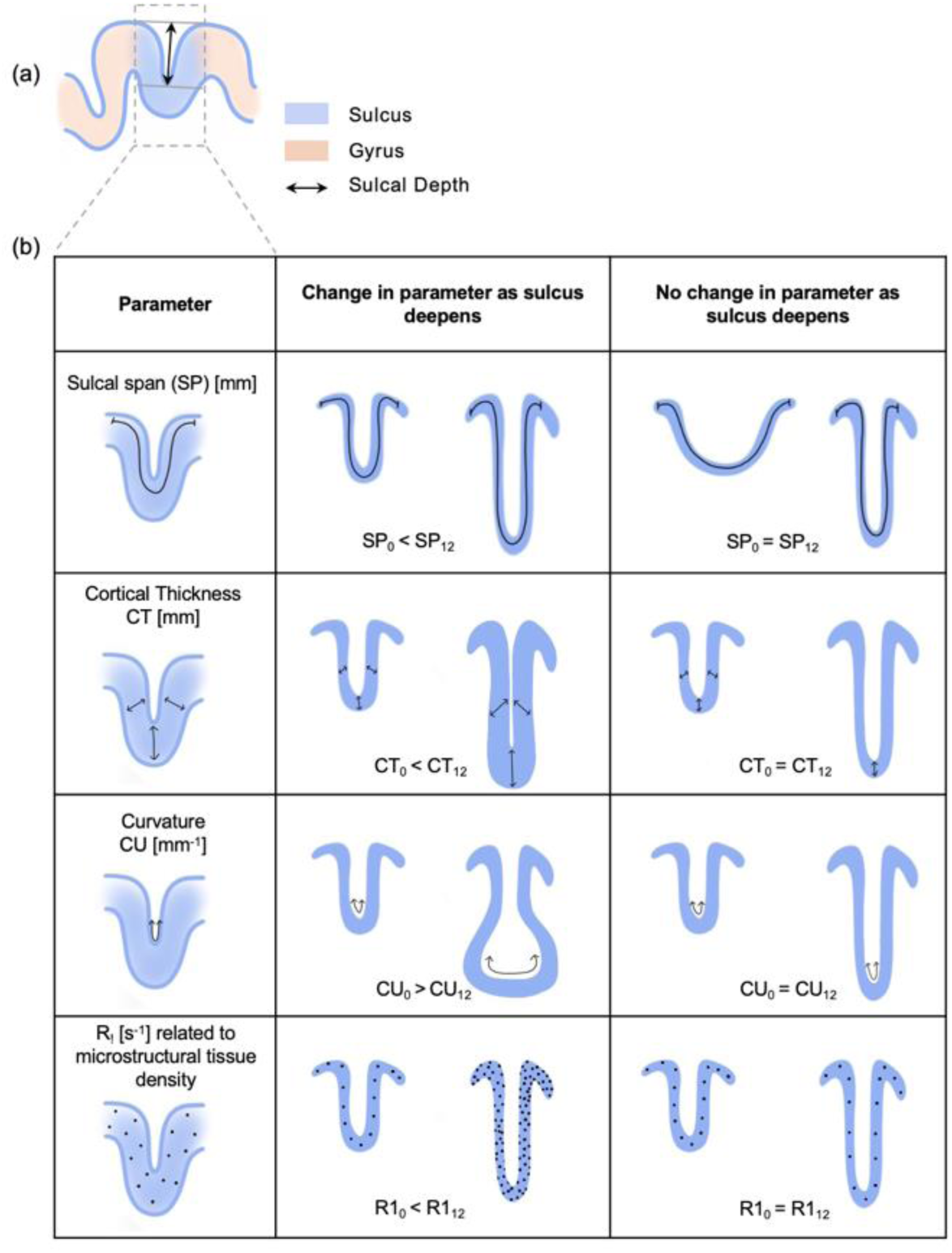
Schematic illustrating possible relationships between sulcal deepening and changes in macrostructural and microstructural parameters during the first year of life. a) Illustration of a cross section of a sulcus (blue) situated between two gyri (orange), with sulcal depth (SD) indicated by a black arrow. b) Each row represents a potential relationship between sulcal deepening and changes in individual macrostructural and microstructural parameters (from birth to 1 year). *Left column:* The parameter changes as the sulcus deepens; *Right column:* The parameter remains unchanged despite sulcal deepening. *Row 1*: Sulcal span (SP). *Row 2:* Cortical thickness (CT). *Row 3:* Cortical curvature (CU). *Row 4:* Tissue relaxation rate (R_1_). Higher R_1_ values correspond to denser cortical tissue (represented by dots).

To test these hypotheses, we conducted longitudinal and cross-sectional anatomical MRI (T_1_/T_2_ weighted) and qMRI in 0 to one-year-old infants during natural sleep. Here, we examine the development of 12 sulci that emerge in utero between the 16th-31st gestational weeks^1^ (**Fig. 2a,b**): calcarine sulcus, parieto-occipital sulcus, insular sulcus, central sulcus, collateral sulcus, superior temporal sulcus, superior frontal sulcus, inferior parietal sulcus, lateral occipital sulcus, inferior frontal sulcus, occipital temporal sulcus, and inferior temporal sulcus. From anatomical MRIs, we estimated sulcal depth [mm], sulcal span [mm], cortical thickness [mm], and cortical curvature [mm^-1^], and from qMRI, we estimated R_1_ [s^-1^] in cortex (**Supplementary Fig. 1** and **Methods**). We examined the development of each of these macroanatomical and microstructural features and their relation to sulcal depth.

**Figure 2.**
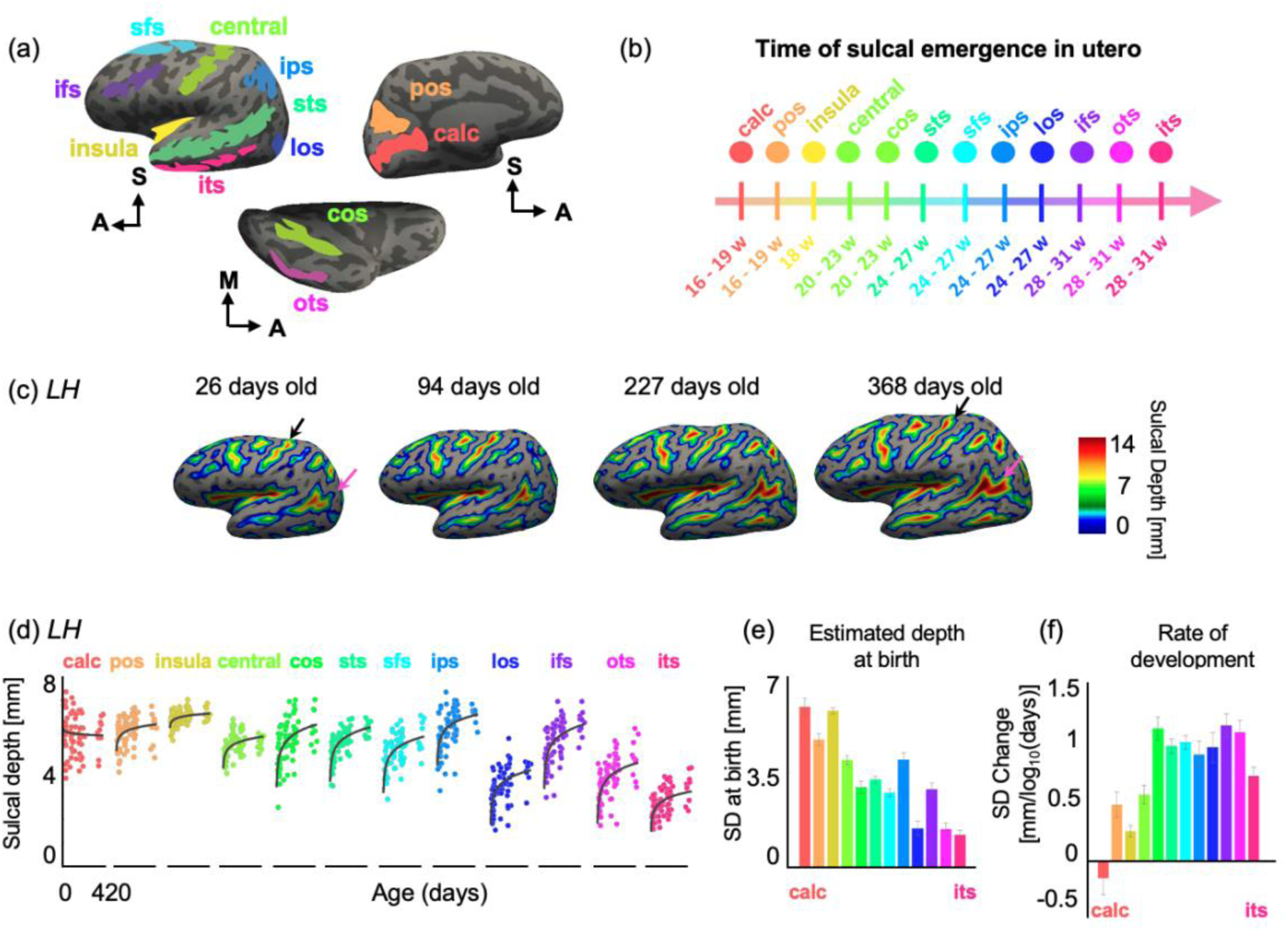
Sulcal folds undergo significant deepening during the first year of postnatal life. (a) 12 primary sulci of interest shown on the cortical surface of an infant. Each color represents a sulcus. (b) Timeline of sulcal emergence in utero during gestation, from Chi et al., (1977). w: week. (c) Sulcal depth (SD in mm) displayed on an inflated cortical surface of an example infant scanned four times over one year. *Left to right:* SD at 26 days (newborn), 94 days (∼3 months), 227 days (∼6 months), and 368 days (∼12 months) (*warmer colors:* higher SD). (d) SD increases logarithmically with age in the first year of life in all sulci except for the calcarine. *Each dot*: mean SD per infant. *Line:* model fit. (e) Estimates of SD at birth (intercept) and (f) rate of development of SD (slope) calculated using LMM relating mean SD versus log_10_ (participant’s age in days) per sulcus. *Error bars:* standard error on estimates of slopes and intercept. Data shown are from the left hemisphere (LH); Right hemisphere data in **Supplementary Fig. 3**. *Abbreviations: calc:* calcarine; *pos:* parieto-occipital sulcus; *insula*: insular sulcus; *central*: central sulcus; *cos*: collateral sulcus; *sts:* superior temporal sulcus; *sfs:* superior frontal sulcus; *ips*: inferior parietal sulcus; *los:* lateral occipital sulcus, *ifs:* inferior frontal sulcus; *ots:* occipital temporal sulcus; *its:* inferior temporal sulcus. Note: the newborn infant (26 days old) shown here is part of the larger infant data set. However, this timepoint was excluded for the remaining analyses due to unusable qMRI data.

## Results

Sixty-one full-term (N_female_ = 24) were recruited to participate in the study. For quality assurance, we (i) monitored each infant’s motion via an infrared camera during each scan, (ii) assessed the quality of brain images, and (iii) repeated scans with motion artifacts. We collected usable data in 79 sessions from 43 infants (N_female_ = 18). 24 of them were scanned longitudinally (**Supplementary Table 1** for demographics; **Methods** for exclusionary criteria). We report data from four timepoints: newborns (*N_sessions_* = 27, N_female_ = 10, mean age ± SD: 29.14 ± 9.92 days), 3-month-olds (*N_sessions_* = 20, N_female=_ 11, 106.35 ± 20.09 days), 6-month-olds (*N_sessions_* = 22, N_female_ =12, 188.45 ± 15.34 days), and 1-year-olds (*N_sessions_*= 10, N_female_ = 3, 385.80 ± 17.60 days).

To test sulcal development, we identified 12 long-length sulci in each infant and timepoint across the brain (**Figs. 2a,b**). These sulci were delineated in the adult average FreeSurfer brain and projected to each infant’s cortical surface using cortex-based alignment^46^ (**Supplementary Fig. 2a,** example infant across four time points). We tested the reliability of automated sulcal fold identification by evaluating the accuracy of automated versus hand-drawn sulci in 10 randomly selected infants across the four timepoints. Dice coefficients (DC) measuring the correspondence between the hand drawn and automated sulcal delineations (DC LH: 0.691 ± 0.14; DC RH: 0.687 ± 0.13) were comparable to DC in our prior work^40^, which compared automated identification of sulci in adult vs. infants brains (DC_adults_: 0.67 ± .05; DC_infants_: 0.66 ± 0.10). This indicates that despite significant brain volume changes during the first year of life, cortex-based alignment using adult brain templates can be used to identify sulci in individual infants’ brains (**Methods** and **Supplementary Fig. 2b**). From these delineations, we calculated mean sulcal depth, mean sulcal span, mean thickness, mean curvature, and mean R_1_ for each sulcus.

### Sulcal depth development is inverse to order of emergence in utero

We first asked if mean SD of the 12 sulci distributed across cortex changes from birth to one year. Visualizing SD maps of the same infant across four timepoints (newborn, 3 months, 6 months, and 12 months) reveals that sulci deepen from birth to one year (**Fig. 2c**). For example, superior temporal sulcus (pink arrow, **Fig. 2c**) and central sulcus (black arrow, **Fig. 2c**) are shallower at 26 days than at 368 days, suggesting increasing depth with age.

To quantify this development, we plotted the mean SD as a function of participants’ age in days and ordered the sulci by their time of emergence in utero (**Fig. 2d)**. In general, SD increases over the first year of life and more rapidly between 0 to 6 months than 6 to 12 months (left hemisphere, LH): **Fig. 2d**, right hemisphere, (RH): **Supplementary Fig. 3a**). Sulcal depth increases on average by ∼21% from birth to 1 year, with an average SD of 4.45 ± 1.30 mm (mean ± standard deviation) in newborns, compared to 5.39 ± 1.07 mm in 1-year-olds. To statistically quantify this development and account for cross-sectional and longitudinal data, we used linear mixed models (*SD ∼ log10(age)*sulcus+(1|infant),* **Methods**). We find that SD significantly varies across sulci in the left hemisphere (main effect of sulcus: LH: *β*= -0.405, SE = 0.04, t_944_ = -9.87, *p*=6.38x10^-22^, 95% CI = [-0.49 -0.32]) and that SD differentially develops across sulci (age by sulcus interaction: LH: *β*= 0.09, SE = 0.02, t_944_ = 4.42, *p*=1.09x10^-05^, 95% CI = [0.05 0.13]) (full statistics for both hemispheres in **Supplementary Table 2**).

Because we found a significant interaction between sulcus and age, we used LMMs to estimate SD at birth and the rate of development of SD per sulcus. Rate of SD change revealed an inverse pattern of development relative to the order of emergence in utero, with earlier emerging sulci being deeper at birth (LH: **Fig. 2e**, RH: **Supplementary Fig. 3b**) but exhibiting slower rates of postnatal deepening compared to later emerging sulci (LH: **Fig. 2f**, RH: **Supplementary Fig. 3c**). Calcarine and insula (emerging first at 16^th^-18^th^ gestational weeks) are the deepest at birth (calcarine = 5.93 ± 0.30 mm; insula = 5.78 ± 0.11 mm) whereas occipital-temporal sulcus (OTS) and inferior temporal sulcus (ITS, emerging at 28^th^-31^st^ gestational weeks) are shallower at birth (OTS = 1.42 ± 0.22 mm; ITS = 1.21 ± 0.17 mm)) (LH: **Fig. 2e**, RH: **Supplementary Fig. 3b,** full statistics per sulcus: **Supplementary Table 3**). At the same time, the calcarine and insula exhibit a non-significant or slower (calcarine: -0.15 ± 0.14 mm/log_10_ (age in days); mm/insula: 0.26 ± 0.05 mm/log_10_ (age in days)) rate of deepening postnatally than later emerging sulci such as the OTS and ITS which deepen more rapidly (OTS: 1.13 ± 0.10 mm/log_10_ (age in days); ITS: 0.75 ± 0.08 mm/log_10_ (age in days)). Hence, later emerging sulci are shallower at birth, but deepen more rapidly from birth to one year.

Having established that sulcal deepening is linked to the timing of emergence in utero, we next examined whether other macrostructural properties—sulcal span (SP), cortical thickness (CT), and curvature (CU)—develop concurrently with sulcal depth (SD) during the first year of life. Using the same approach as SD, we applied linear mixed models (LMMs) to estimate the birth values and rates of change for each parameter, allowing us to assess whether they follow similar developmental trajectories or exhibit distinct patterns across sulci.

### Sulci widen with age in the first year of life

The sulcal span of all 12 sulci increases from birth to 12 months. This increase in span, like that for SD, is faster between birth to 6 months, than from 6 to 12 months (LH: **Fig. 3a**, RH: **Supplementary Fig. 3d**). We observed an average increase of 42% in span from newborns (24.20 ± 7.39 mm) to one-year-olds (34.31 ± 10.97 mm). We find that sulcal span significantly varies with age (main effect of age: LH: *β*=8.44, SE=1.36, t_944_=6.20, *p*=8.06x10^-10^, 95% CI =[5.77 11.10]) and across sulci (main effect of sulcus: LH: *β*=-0.77, SE=0.37, t_944_=-2.08, *p*=0.037, 95% CI=[-1.5 -0.04]) (full statistics in **Supplementary Table 2**). The estimates of sulcal span at birth showed a pattern like that of SD, whereby earlier emerging sulci like the calcarine sulcus are deeper at birth than later emerging folds like ITS. However, unlike that for SD, the development of sulcal span does not follow a developmental pattern related to the time of emergence (LH: **Figs. 3b-c**, RH: **Supplementary Figs. 3e-f,** full statistics per sulcus: **Supplementary Table 4**). These data indicate that sulcal span increases significantly during the first year of life, reflecting ongoing cortical expansion. However, unlike sulcal depth, the rate of span expansion does not systematically follow the timing of sulcal emergence in utero.

**Figure 3.**
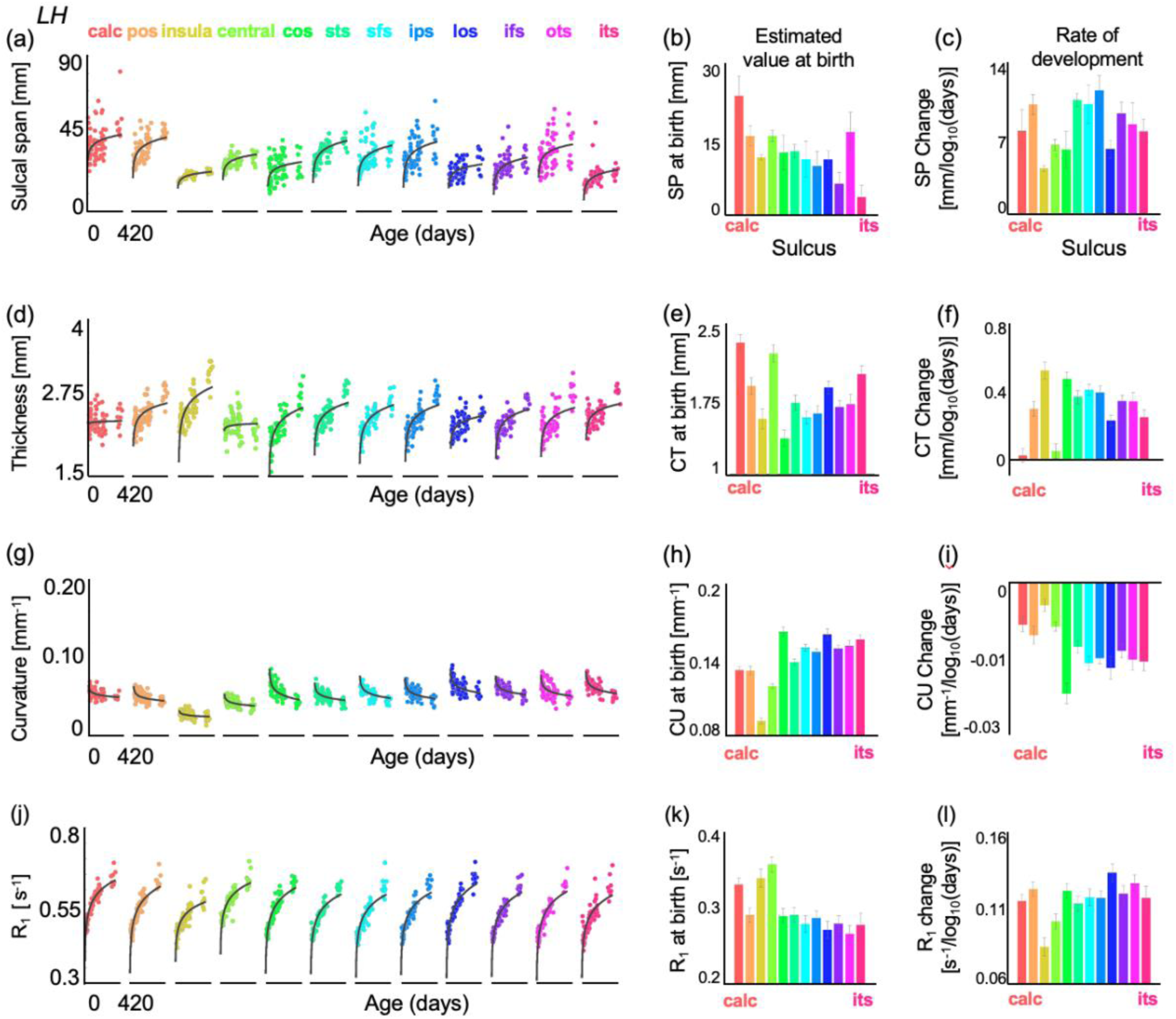
Macroanatomical and microstructural changes in sulci during the first year of postnatal life. (a) Sulcal span (SP [mm]) increases logarithmically with age across all sulci in the first year of life. *Each dot*: mean SP per infant. *Line:* model fit. (b-c) Estimates of SP at birth (intercept) and rate of development (slope), calculated using a LMM relating mean SP to log_10_ (participant’s age in days). *Error bars:* standard error on estimates of slopes and intercept. (d-l) Same format as (a-c), shown for cortical thickness (CT [mm]), curvature (CU [mm^-1^]), and cortical microstructure (R_1_ [s^-1^]). Like SP, CT and R_1_ increase with age, while CU decreases. All data shown are from the left hemisphere (LH). Right hemisphere data are in **Supplementary Fig. 3**.

### Cortex thickens in sulcal folds from birth to one year of life

Next, we quantified the development of cortical thickness (CT) of the sulci during the first year of life. Like depth and span, CT also significantly increases from birth (2.27 ± 0.169 mm to one year (2.74 ± 0.26 mm, ∼21% increase) following a logarithmic profile (LH: **Fig. 3d**, RH: **Supplementary Fig. 3g**). Cortical thickening is faster earlier on, between birth to about 6 months of life, than later between 6 to 12 months (LMM: *CT* ∼ *log*10(*age of infant*) ∗ *sulcus* + (1|*infant*); main effect of age: LH: *β*= 0.25, SE = 0.03, t_944_ = 8.19, *p*=8.30x10^-16^, 95% CI =[0.19 0.31]). CT also varies across sulci (main effect of sulcus: LH: *β*= -0.02, SE = 0.008, t_944_ = -2.72, *p*=0.006, 95% CI = [- 0.037 0.0061]) and shows sulcus-specific developmental trajectories (age by sulcus interaction: LH: *β*= 0.0095, SE = 0.004, t_944_ = 2.38, *p*=0.017, 95% CI = [0.0016 0.017]) (full statistics in **Supplementary Table 2**). Across individual sulci, cortical thickness significantly increases in 10 out of the 12 sulci (exceptions are the calcarine and central sulcus, full statistics per sulcus: **Supplementary Table 5**). However, the rate of cortical thickening does not follow a clear pattern based on time of emergence in utero. (LH: **Figs. 3e-f**, RH: **Supplementary Figs. 3h-i**). Instead, sulci that are thicker at birth, such as the calcarine and central sulcus, show minimal thickening, whereas sulci that are thinner at birth, such as the collateral sulcus and insula, exhibit substantial thickening during the first year. These data suggest that postnatal cortical thickening follows distinct maturational trajectories that are more closely tied to cortical thickness at birth than to the timing of sulcal emergence in utero.

### Curvature of sulcal folds decreases and becomes less concave in the first year of life

Next, we examined whether sulcal curvature (CU) changes during the first year of life. Unlike SD, SP, and CT, which increase with age, CU decreases. On average, CU declines by 14% from birth (0.12 ± 0.01 mm^-1^) to one year of age (0.11 ± 0.01 mm^-1^) (LH: **Fig. 3g**, RH: **Supplementary Fig. 3j**). Curvature significantly decreases following a logarithmic trajectory (LMM: *CU* ∼ *log*10(*age of infant*) ∗ *sulcus* + (1|*infant*); main effect of age: LH: *β*= -0.0095, SE = 0.0019, t_944_ = -4.82, *p*=1.64x10^-06^, 95% CI =[-0.013 -0.0056]). Curvature also varies across sulci (main effect of sulcus: LH: *β*= 0.0035, SE = 0.00054, t_944_ = 6.53, *p*=1.03x10^-10^, 95% CI = [0.0024 0.0045]) and develops differentially across sulci (age by sulcus interaction: LH: *β*= -0.00067, SE =0.00027, t_944_ = -2.49, *p*=0.013, 95% CI = [-0.0012 -0.00014]) (full statistics in **Supplementary Table 2**). Some sulci, such as the calcarine and insula, show minimal curvature change, while others, like the STS and ITS, become markedly less curved over the first year (LH: **Figs. 3h-i**, RH: **Supplementary Figs. 3k-l,** full statistics per sulcus: **Supplementary Table 6**). Developmental decreases in mean curvature may result from sulcal widening at the fundus, increased cortical expansion along the sulcal walls, or a combination of both. Notably, curvature changes are not uniform across sulci, suggesting that different sulci undergo distinct developmental trajectories postnatally.

### Sulci undergo extensive microstructural tissue growth in gray matter

Next, we investigated if microstructural tissue properties of gray matter of these 12 sulci, measured using quantitative R_1_ [s^-1^] develop with age. We find that R_1_ increases with age (LH: **Fig. 3j**, RH: **Supplementary Fig. 3m**) and this development is faster earlier on, between birth to about 6 months of life, than later between 6 to 12 months. R_1_ increases on average by ∼33% from newborns (0.46 ± 0.02 s^-1^) to one-year-olds (0.61 ± 0.03 s^-1^) (LMM: *R*1 ∼ *log*10(*age of infant*) ∗ *sulcus* + (1|*infant*); main effect of age: LH: *β*=0.104, SE = 0.0039, t_944_ = 26.59, *p*=9.73x10^-117^, 95% CI =[0.096 0.11]). R_1_ also varies across sulci (main effect of sulcus: LH: *β*= -0.0058, SE = 0.0009, t_944_ = -5.91, *p*=4.79x10^-09^, 95% CI = [-0.008 -0.004]), and finally, R_1_ differentially develops across sulci (age by sulcus interaction: LH: *β*= 0.0015, SE =0.0005, t_944_ =3.11, *p*=0.0019, 95% CI = [0.00056 0.0025]) (full statistics for both hemispheres in **Supplementary Table 2**). R_1_ development is less orderly by time of emergence (LH: **Figs. 3k-l and** RH: **Supplementary Figs. n-o;** full statistics per sulcus: **Supplementary Table 7**). These results reveal that sulci become microstructurally denser over the first year of life, indicating extensive tissue growth.

### Sulcal depth development can be predicted by a linear combination of sulcal span, thickness, curvature, and R_1_

Thus far, our data indicate that macrostructural and microstructural parameters undergo distinct age-related changes across the 12 sulcal folds. As a sulcus deepens, its span widens, cortex thickens, curvature decreases, and microstructural tissue density increases. **Figure 4a** summarizes these developmental changes using a sample infant’s left collateral sulcus from birth to one year of age. However, the magnitude and trajectory of these changes vary across sulci, as shown in **Figures 2** and **3**. Given these findings, an open question is whether sulcal deepening is systematically driven by concurrent macrostructural and microstructural developments.

**Figure 4.**
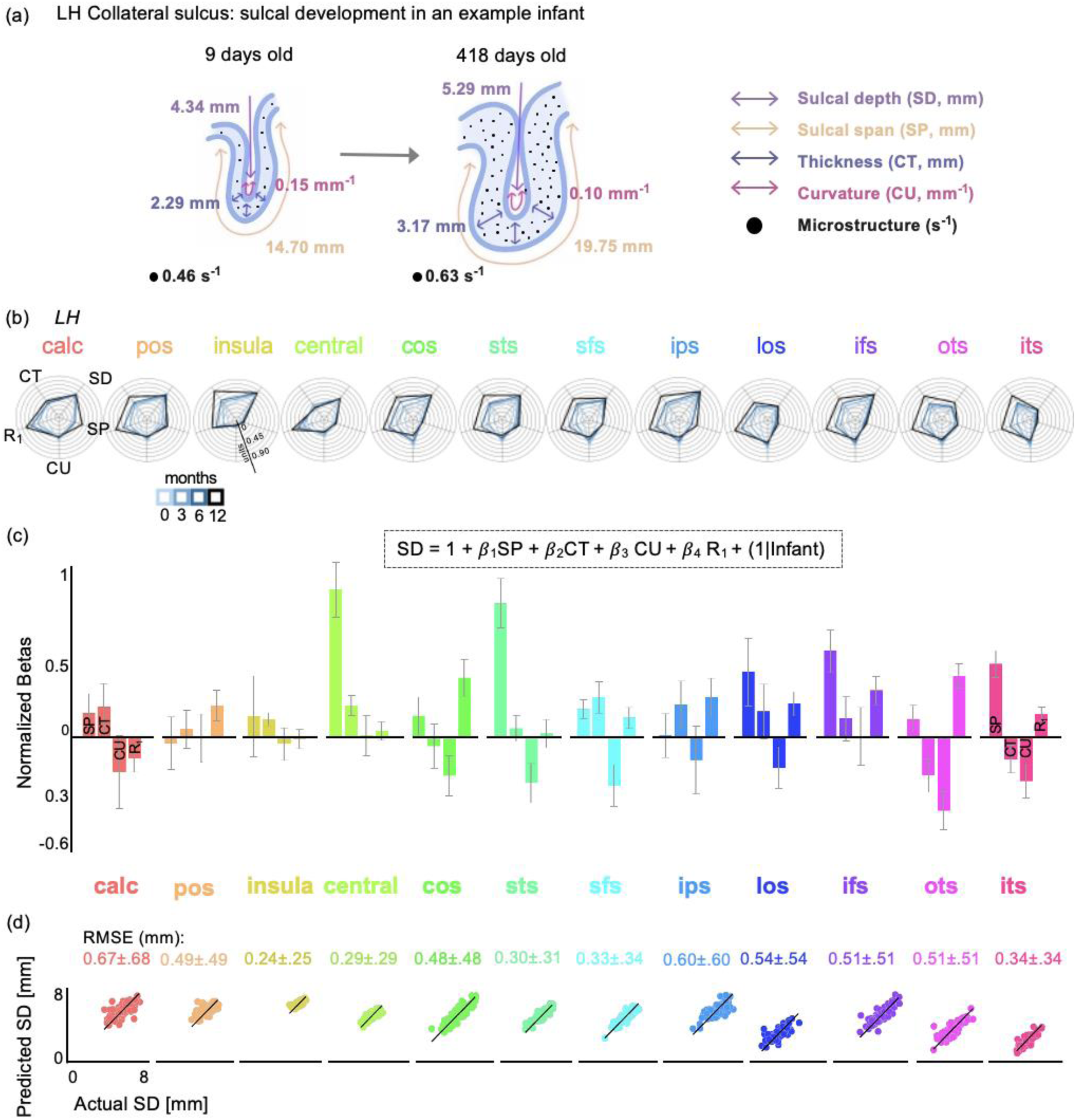
SD development can be predicted by a linear combination of SP, CT, CU, and R_1_. (a) Summary schematic showing macroanatomical and microstructural changes occurring in a sample infant’s left collateral sulcus from birth to one year. (b) Polar plots representing normalized anatomical and microstructural properties per sulcus. Four solid lines (light blue – 0 months to dark blue – 12 months) represent participants’ age group. Each concentric circle corresponds to the normalized units to allow the metrics to be plotted with respect to each other. (c) Normalized beta values from a linear model relating SD to SP, CT, CU, and R_1_. (d) Scatterplots displaying predicted mean depth [mm] in left out participants using the model parameters in (c) versus actual mean depth [mm] per sulcus. Each dot is a participant. Root means square error (RMSE [mm]) per model is reported per sulcus. *LH*: left hemisphere. RH data in **Supplementary Figure 4**.

First, to visualize how the individual parameters develop and vary with respect to each other across the sulci, we generated polar plots visualizing the normalized metrics in the 12 sulci across four age groups (0, 3, 6, and 12 months) (LH: **Fig. 4b**, RH: **Supplementary Fig. 4a**). As parameters have different units and ranges, we normalized all metrics to visualization and analysis (see **Methods**). Polar plots show distinct developmental fingerprints across sulci. For instance, while both CoS and STS show substantial increases in tissue density (R_1_), the STS shows proportionally greater expansion in span. These differences suggest that the relationships between sulcal depth and other cortical properties may not be uniform across sulci. To quantify these relationships, we first used a global LMM to test if the relation between sulcal depth and sulcal span, cortical thickness, curvature, and microstructural tissue density varies across sulci. Because all parameters exhibited significant age-related changes, we included age in the model to isolate the independent effects of macrostructural and microstructural properties on sulcal depth. Thus, we examine the relation between SD and the other parameters as follows: *SD* ∼ 1 + (*Age* + *SP* + *CT* + *CU* + *R*1) ∗ *sulcus* + (1|*infant*). The model used normalized parameters, consistent with the visualizations above.

As expected, across all sulci, sulcal depth varies with age (main effect of age, *β* = 0.755; SE=0.27; t_936=_> 2.79, ps<0.0054; no significant interactions). But consistent with the patterns observed in the polar plots, the relationship between depth and macrostructural (span, thickness, curvature) and microstructural (R_1_) parameters significantly varies across sulci (full statistics for both LH and RH in **Supplementary Table 8;** LH: significant interaction between SP and sulcus: *β* = 0.00353, SE = 0.0001, t_936_ = 3.94, *p* = 8.8 x 10^-05^, 95% CI = [0 0.01]); significant interaction between CT and sulcus: *β* = -0.215, SE = 0.04, t_936_ = -5.14, *p*=3.4x 10^-07^, 95% CI = [-0.3 -0.13]); significant interaction between CU and sulcus: *β* = -4.36, SE = 0.74, t_936_ = -5.9, *p* = 4.96 x 10^-05^, 95% CI = [- 5.82 -2.91]; significant interaction between R_1_ and sulcus: *β* = 0.836, SE = 0.29, t_936_ = 2.93, *p* = 0.00342, 95% CI = [0.28 1.4]). These interactions indicate that sulcal deepening is not governed by a single uniform process but emerges from distinct structural factors that differ across folds.

Thus, we next examined these relationships within individual sulci. For each sulcus, we ran a separate LMM: (*SD* ∼ 1 + (*SP* + *CT* + *R*1 + *CU*) + (1|*infant*)), to estimate beta coefficients for each parameter. Overall, we found positive beta coefficients for SP and CT, whereas beta coefficients for curvature and R_1_ exhibited both positive and negative values depending on the sulcus (LH: **Fig. 4c**; RH**: Supplementary Fig. 4b; Supplementary Table 9** for beta values per parameter and hemisphere). As a control, we also ran an LMM per sulcus adding age as factor: (*SD* ∼ 1 + (*Age* + *SP* + *CT* + *R*1 + *CU*) + (1|*infant*)), finding an overall similar pattern of results (**Supplementary Fig. 5**).

To assess the robustness of our model, we tested whether sulcal depth can be predicted in new participants from these parameters using a leave-one-out cross-validation approach. Unlike the previous analyses, this model was trained and evaluated using raw (non-normalized) values to preserve interpretability in millimeters. Specifically, we trained the model on n-1 infants and used it to predict sulcal depth in the left-out infant. Results revealed that mean sulcal depth can be reliably predicted using our model, with an average root-mean-square-error (RMSE) of less than 1mm (LH: **Fig. 4d**; RH**: Supplementary Fig. 4c**). However, prediction accuracy varied across sulci, with the insula yielding the lowest prediction error (RMSE = 0.25 mm) and the calcarine yielding the highest (RMSE = 0.67 mm). Overall, our results highlight that multiple morphological (span, thickness, curvature) and microanatomical (R_1_) developments predict sulcal depth during the first year of life, even as their weights vary across sulci.

### Strong relationship between depth, curvature, and microstructure along a sulcal fold

Our findings thus far suggest that mean sulcal depth is systematically related to macro- and microstructural properties of cortex. However, mean sulcal depth measures inherently average data across different portions of a sulcus, e.g., over the fundus and sulcal walls, which may obscure meaningful fine-grained variation. Prior work has shown that functional properties often vary along the sulcal fold, with stronger functional coupling in the fundus versus the walls^13–15,47–51^. Motivated by prior work, we asked whether sulcal depth along the sulcus is spatially coupled with micro and macrostructural properties of cortex during infancy. Specifically, we examined whether depth varies systematically along each sulcal fold by quantifying the relationship between local sulcal depth, thickness, curvature, and R_1_. In this analysis, we did not include sulcal span because our metric of sulcal span is a single measure per sulcus and does not capture localized variations in span along the sulcus.

We first visualize how depth, cortical thickness, curvature, and R_1_ vary along each sulcus in individual infants. As an example, **Fig. 5a** shows the left superior temporal sulcus (STS) in an example one-year-old infant. Notably, the STS has two distinct deep regions (red peaks in **Fig. 5a**-left). Comparing these depth variations to curvature, R_1_, and thickness along the sulcus (**Figs. 5a,b**) revealed that (1) curvature follows the depth pattern, with greater curvature in the two deepest regions of the sulcus than in shallower areas, (2) R_1_ is higher in deeper regions of the sulcus, and (3) cortical thickness exhibits a weaker relationship to sulcal depth. We also provide an example for the right CoS in a different one-year-old infant (**Supplementary Fig. 6**), which illustrates a strong relationship between depth and both CU and R_1_ along the CoS.

**Figure 5.**
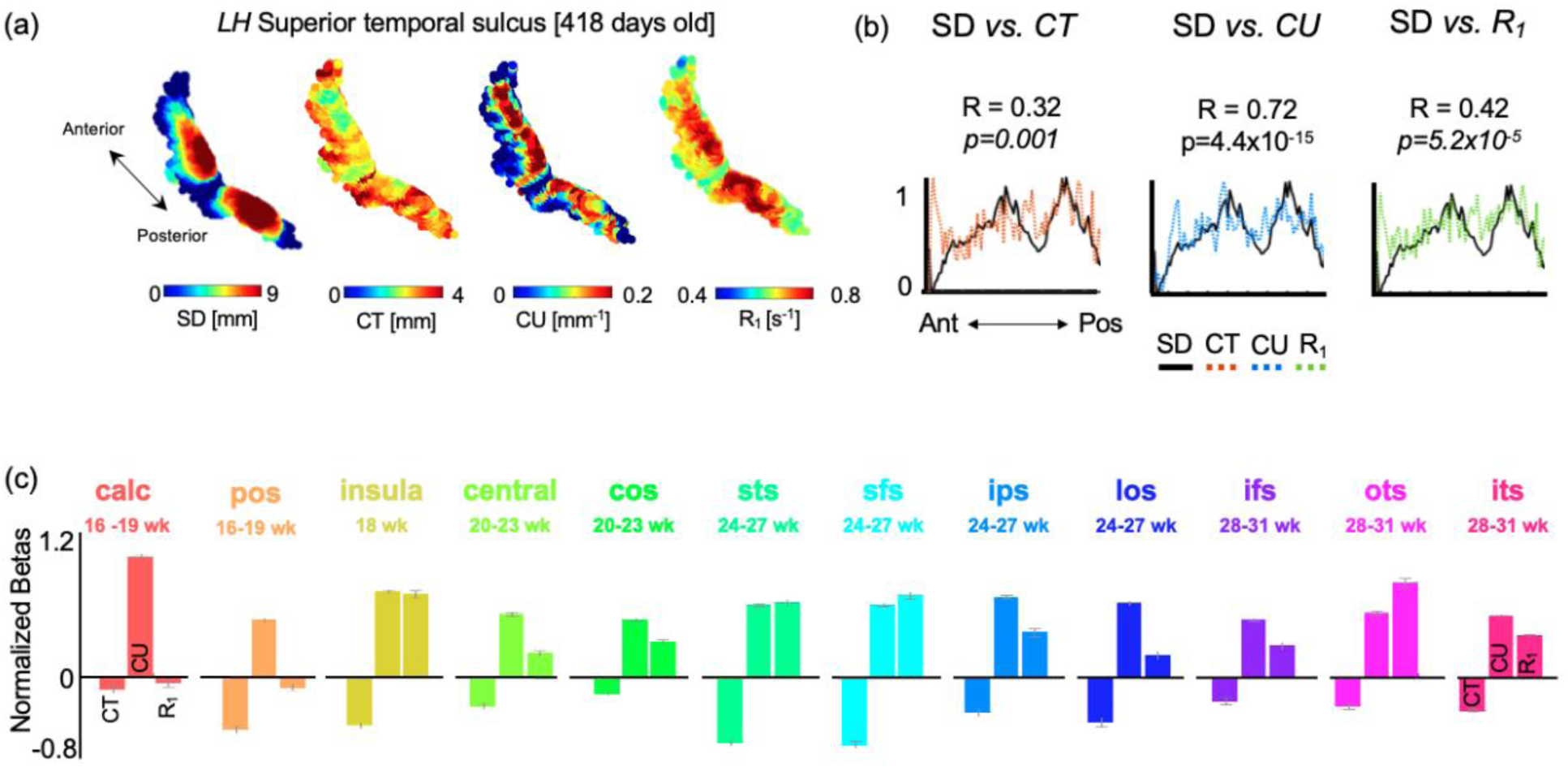
Strong coupling between depth versus macro- and microstructural properties along the sulcal fold. (a) Three-dimensional maps of left superior temporal sulcus (STS) of a sample one-year-old (418 days) showing depth, curvature, R_1_, and CT along the sulcus; warmer colors represent larger values. (b) Normalized depth (solid black lines), CT (orange dotted lines), CU (blue dotted lines), and R_1_ (green dotted lines) values as a function of distance along the anterior-posterior axis of the STS. Data show the coupling between depth and each parameter. Mean and standard deviation of the correlation (R) and significance values (p) are provided in each subplot. (c) Bar plots showing normalized beta values from a linear model relating SD to CT, CU, and R_1_ along the length of each sulcus. *LH*: left hemisphere. **Supplementary Fig. 6** shows right hemisphere data.

To quantify these fine-grained relationships, we used an LMM (*SD* ∼ 1 + (*CT* + *CU* + *R*1) ∗ *sulcus* + (1|*infant*), see **Methods**). Results revealed that local sulcal depth is significantly related to local CT, CU, and R_1_ (32.69<|Ts|<102.75; ps<8.61x10^-5^) and that the relationships significantly vary across sulci (LH: significant interaction between CT and sulcus *β* = 0.01, SE = 0.002, t_45486_ = 4.63, *p* = 3.64x10^-06^, 95% CI = [0.005 0.013]); significant interaction between CU and sulcus: *β* = -0.05, SE = 0.002, t_45486_ = -30.08, *p* = 7.34x10^-197^, 95% CI = [-0.05 -0.04]; significant interaction between R_1_ and sulcus: *β* = 0.05, SE = 0.002, t_45486_ = 24.14, *p* = 6.66x10^-128^, 95% CI = [0.05 0.06]; statistics for both hemispheres: **Supplementary Table 10).** Given these interactions, we next estimated the relationship between local sulcal depth and CT, CU and R_1_ per sulcus using a separate LMM: *SD* ∼ 1 + (*CT* + *CU* + *R*1) + (1|*infant*). We found that deeper points along sulci tended to have higher curvature and R_1_, and lower cortical thickness, except for the calcarine and POS, where R_1_ coefficients were close to zero or negative (LH: **Fig. 5c**; RH**: Supplementary Fig. 6c; Supplementary Table 11** for beta values per parameter and hemisphere). Notably, this fine-grained analysis showed a positive relationship between local sulcal depth and local curvature, in contrast to the prior analysis, which found a negative relationship between their mean values (**Fig. 4c; Supplementary Table 9**). This difference is not due to model differences, as mean sulcal depth is negatively related to mean curvature even when sulcal span is excluded from the model. Rather, this shift in direction likely reflects the different levels of analysis. While mean curvature averages across the entire sulcus—including both the fundus and sulcal walls—local analysis preserves anatomically meaningful variations along the fold. These findings underscore the importance of within-sulcus analyses for capturing fine-grained structural features that may be obscured by averaging.

Overall, these findings show that deeper points along sulci tend to have higher curvature, higher microstructural density tissue, and lower thickness. Higher microstructural tissue density and curvature in deeper portions of sulci (e.g., fundi) suggests that sulcal fundi may undergo distinct tissue development compared to sulcal walls. Moreover, the negative relationship between depth and thickness is consistent with prior work showing that thickness declines linearly with depth^52^.

## Discussion

Our findings reveal a dynamic interplay between sulcal macro- and microstructural tissue density during infant cortical development. Using both cross-sectional and longitudinal data, we provide the first systematic analysis of anatomical and microstructural changes in individual sulcal folds during the first year of postnatal life. First, our work reveals a fine-grained, hierarchical, and sulcus-specific developmental pattern whereby early emerging sulci are deeper at birth but deepen more slowly postnatally compared to later emerging sulci. Second, concurrent with sulcal deepening, we observe increases in sulcal span, cortical thickness, and microstructural tissue density, accompanied by reductions in cortical curvature. Third, our results reveal distinct yet strong relationships between depth, curvature, thickness, and microstructural tissue density along the length of sulci, highlighting local structural coupling within sulcal folds. Finally, by modeling sulcal depth as a weighted combination of multiple anatomical parameters, we find that sulcal deepening does not emerge from a singular mechanism but instead reflects the interplay of multiple interacting macroanatomical and microstructural processes.

### Sulcal depth development postnatally is anchored to time of emergence in utero

Our results highlight a consistent yet heterogeneous trajectory of sulcal deepening after birth. Here we find sulcus-specific developmental patterns shaped by the timing of sulcal emergence in utero. Specifically, early emerging sulci (e.g., calcarine and insula) are deeper at birth but show minimal postnatal deepening, while later-emerging sulci (e.g., occipital temporal and inferior temporal sulcus) are shallower at birth but deepen more rapidly postnatally. This pattern aligns with prenatal imaging studies^53,54^ that describe a sequential wave of sulcal maturation from central to more lateral and anterior regions. This progression may reflect early fetal cytoarchitectonic patterning governed by a genetically encoded proto-map established early in fetal development^20,54,55^. Our findings support the view that sulcal maturation during infancy is not random but rather follows an organized spatial and temporal trajectory, established prenatally.

### Macrostructural and microstructural changes co-occur with sulcal deepening

Alongside sulcal deepening, we found that mean sulcal span, mean cortical thickness, and mean R_1_ significantly increase during the first year of life, while mean curvature decreases. These coordinated changes indicate that sulci become wider, thicker, denser, and less concave from birth to 1 year. However, in contrast to sulcal depth, the development of other features does not systematically vary with gestational emergence. This dissociation suggests that sulcal depth may be more directly shaped by early prenatal influences, while other macrostructural and microstructural properties undergo differential developments during infancy. Our findings also align with several hypothesized mechanisms of sulcal deepening proposed in **Figure 1**. The simultaneous increases in sulcal span and depth are consistent with cortical expansion in the tangential direction^44^ (parallel to the cortical surface), which may stretch folds laterally and drive them deeper into white matter due to spatial constraints imposed by the skull^12^. The increase in mean sulcal depth and mean cortical thickness aligns with radial expansion^5^ (perpendicular to the cortical surface). The observed decrease in mean curvature suggests that sulci may become less concave (vase-shaped) during this time of development, possibly due to widening at the fundus and/or lengthening or stretching along sulcal walls. Finally, increasing cortical R_1_ values point to extensive tissue growth, which may further expand cortex and/or increase its stiffness, thereby contributing to internal folding pressures that push sulci deeper into white matter. Our novel finding that depth is positively associated with R_1_—both on average and locally—suggests that changing tissue density may exert mechanical forces that contribute to sulcal deepening. Together, these results underscore that an array of differentially developing macroanatomical and microstructural processes contribute to sulcal deepening during the first year of human life.

### Implications for biomechanical models of cortical folding

Cortical folding is often modeled as the mechanical outcome of differential expansion and structural constraints, incorporating metrics like cortical thickness, surface area, and gray-white matter elasticity^4,21,41,56^. These models have successfully demonstrated how large-scale folding patterns can emerge from physical forces acting on an elastic, largely homogeneous cortical sheet. However, general folding mechanisms alone are insufficient to explain the sulcus-specific developmental variability that we observe across spatial location and gestational timing. Our findings, along with insights from large-scale developmental datasets^57–59^, highlight the need for biomechanical models to incorporate sulcus-specific heterogeneity. These empirical patterns offer critical constraints for building more nuanced, data-driven models of cortical folding. Additionally, most existing simulations—including gel-based physical models—do not account for fine-grained anatomical variation within sulci, which limits the ability to fully explain how folding develops. Our finding that the deepest portions of sulci (fundi) consistently exhibit higher microstructural density (R_1_) and curvature suggests that localized maturational and mechanical forces may differentially influence the development of sulcal fundi compared to their surrounding walls. Future models that implement localized variation in tissue density (e.g., modeling higher R_1_ as increased stiffness or reduced elasticity) could provide new insight into how mechanical forces within the gray matter contribute to cortical folding at finer spatial resolutions.

### Sulcal maturation as a potential biomarker for development of function

While our study focused on structural development, our findings offer insight into how sulcal maturation may relate to the emergence of function during infancy^60^. Sulcus-specific developmental trajectories suggest that the timing of sulcal emergence could influence when and how cortical functions begin to organize. For instance, the superior temporal sulcus, which contains functional regions involved in higher-order processes such as social communication and language, continues to develop structurally throughout infancy and childhood, showing marked changes in depth, curvature, thickness, span, and microstructure. In contrast, sulci like the calcarine, which contains primary visual cortex (V1) responsible for processing basic visual features such as local edges and contrast^61^, appear more structurally mature at birth. These observations raise the possibility that anatomical and functional development unfold in tandem. Although this idea of coupling between structural and functional development aligns with influential findings from animal models^62^, structural developments have yet to be incorporated into computational models of human brain development, which typically model function alone^63–68^.

Our results also suggest that the fundus, or deepest point of a sulcus, may serve as a meaningful biomarker in early development. First, fundi exhibit higher curvature and R_1_ than sulcal walls, suggesting distinct maturational trajectories between these regions at both the macro- and microstructural levels. Second, larger R_1_ increases in fundi than adjacent sulcal walls during infancy suggest that fundi may be localized zones of enhanced microstructure. R_1_ increases may be coupled with increased myelination^33,39,40^, synapse formation^30,62^, and dendritic growth^32^, which in turn may affect cortical function. Indeed, previous studies have shown that functional selectivity is higher in fundi. For example, we found that place-selectivity is higher in the fundus of the collateral sulcus in both children and adults, and this functional-structural coupling strengthens with age^14^. Likewise, deeper regions in the superior temporal sulcus have been linked to language and social processing^13,49,51^, and deeper portions of the central sulcus are associated with motor representations of the hand^48,50^. Together, our findings suggest that future studies should examine the fine-grained relationship between anatomical and functional development during infancy^69–72^ and in particular, the normative development of fundi^18,51^.

### Implications for clinical research

Understanding sulcal development is clinically significant, as deviations from typical folding patterns have been associated with neurodevelopmental disorders. Sulcal depth irregularities are biomarkers of neurodevelopmental disorders such as autism^17,73^, Down syndrome^16^, and schizophrenia^74^. By providing normative developmental trajectories of sulcal deepening, our work establishes a benchmark for identifying early deviations indicative of neurological disorders. Furthermore, the strong association between sulcal depth and microstructural tissue density suggests that disruptions in cortical tissue properties, rather than just altered folding patterns, may contribute to atypical brain development. Additionally, our predictive model accurately generalized sulcal depth estimates to novel infant data, indicating that developmental trajectories are remarkably conserved despite individual variability. This suggests that deviations from expected sulcal deepening patterns could serve as early indicators for identifying at-risk populations.

In conclusion, our findings reveal differential patterns of mean and local sulcal development, each linked to multiple macrostructural and microstructural properties. Together, these patterns offer a novel and promising framework for quantifying and detecting departures from typical neurodevelopmental trajectories. As infants are a highly vulnerable population, the ability to diagnose delayed or atypical development early is critical. The data and methods in our study have important implications for enhancing early detection of neurodevelopmental delays, which may support timely intervention and improved long-term outcomes.

## Methods

### Participants

Sixty-one full-term and healthy infants (N_female_ = 24) were recruited to participate in the study. Forty-three out of sixty-one infants provided usable data. We excluded data from infants that could not fall asleep inside the MRI scanner, which led to excessive motion and unusable data (see additional exclusion criteria in section titled “Expectant parent and infant screening procedure”). Hence, we report longitudinal and cross-sectional data from 43 infants (N_female_ = 18) gathered over 79 scanning sessions, across 4 timepoints: newborns (*N_sessions_* = 27; 10 females; M_age_ ± SD: 29.14 ± 9.92 days), 3 month-olds (*N_sessions_* = 20; 11 females; 106.35 ± 20.09 days), 6 month-olds (*N_sessions_* = 22; 12 females; 188.45 ± 15.34 days), and 1-year-olds (*N_sessions_*= 10; 3 females; 385.80 ± 17.60 days). 24 out of the 43 infants participated in 2 or more timepoints (**Supplementary Table 1**). The participant population was racially and ethnically diverse, reflecting the population of the San Francisco Bay Area, including 1 Hispanic, 5 Asian, 20 Caucasian, and 17 multiracial participants (**Supplementary Table 1**).

### Expectant parent and infant screening procedure

Expectant parents and their infants in our study were recruited from the San Francisco Bay Area using social media platforms. We performed a two-step screening process. First, parents were screened over the phone for eligibility based on exclusionary criteria designed to recruit a sample of typically developing infants. Second, eligible expectant mothers were screened once again after giving birth. Exclusionary criteria were as follows: recreational drug use during pregnancy, significant alcohol use during pregnancy (more than three instances of alcohol consumption per trimester; more than 1 drink per occasion), lifetime diagnosis of autism spectrum disorder or a disorder involving psychosis or mania, taking prescription medications for any of these disorders during pregnancy, insufficient written and spoken English ability to understand the instructions of the study, or learning disabilities that would preclude participation. Exclusionary criteria for infants were preterm birth (<37 gestational weeks), low birthweight (<5 lbs 8 oz), small height (<18 inches), any congenital, genetic, and neurological disorders, visual problems, complications during birth that involved the infant (e.g., NICU stay), history of head trauma, and contraindications for MRI (e.g., metal implants). Study protocols were approved by the Stanford University Internal Review Board on Human Subjects Research. Participants were compensated with 25 dollars per hour for their participation.

### Data Acquisition

All included infants completed multiple scanning protocols to obtain anatomical and quantitative MRI data in a 3T GE scanner at the Center for Cognitive and Neurobiological Imaging at Stanford University. Scanning was done with first level SAR to ensure infants’ safety. Of the 79 scanning sessions, 68 were collected using a Nova 32-channel head coil, and the remaining 11 used a custom 32-channel infant head coil^75^.

Scanning sessions were scheduled in the evenings around the infants’ typical bedtime. Each session lasted between 2.5-5 hours including time to prepare the infant and waiting time for them to fall asleep. Upon arrival, caregivers provided written, informed consent for themselves and their infant to participate in the study. Before entering the MRI suite, both caregiver and infant were checked to ensure that they were metal-free, and caregivers changed the infant into MR-safe cotton onesies and footed pants provided by the researchers. The infant was swaddled with a blanket with their hands to their sides to avoid their hands creating a loop and the researchers inserted soft wax earplugs into the infant’s ears. During sessions involving newborn infants, an MR-safe plastic immobilizer (MedVac, www.supertechx-ray.com) was used to stabilize the infant and their head position. Once the infant was ready for scanning, the caregiver and infant entered the MR suite. The caregiver was instructed to follow their child’s regular sleep routine. When the infant was asleep, the caregiver placed the infant on the scanner bed. Weighted bags were placed at the edges of the bed to prevent any side-to-side movement. Additional pads were also placed around the infant’s head and body to stabilize head position. MRI compatible neonatal noise attenuators (https://newborncare.natus.com/products-services/newborn-care-products/nursery-essentials/minimuffs-neonatal-noise-attenuators) and headphones (https://www.alpinehearingprotection.com/products/muffy-baby) were placed on the infant’s ears, to lower sound transmission. An experimenter stayed inside the MR suite with the infant during the entire scan.

For additional monitoring of the infant’s safety and tracking of the infant’s head motion, an infrared camera was affixed to the head coil and positioned for viewing the infant’s face in the scanner. The researcher operating the scanner monitored the infant via the camera feed, which allowed for the scan to be stopped immediately if the infant showed signs of waking or distress. This setup also allowed tracking the infant’s motion; scans were stopped and repeated if there was excessive head motion. To ensure scan data quality, in addition to real-time monitoring of the infant’s motion via an infrared camera, MR brain image quality was also assessed immediately after acquisition of each sequence and sequences were repeated if necessary.

### Data acquisition parameters and preprocessing

#### Anatomical MRI

T1-weighted and T2-weighted images were acquired and used for tissue segmentation. T1-weighted image acquisition used GE’s BRAVO sequence with TE=2.7ms, TR=6.7ms, echo train length = 1; voxel size = 1 mm^3^; Scan time: ∼3 min. T2-weighted image acquisition used GE’s CUBE sequence with TE=122ms, TR = 3650 ms; echo train length = 120; voxel size = 1 mm^3^; FOV = 20.5 cm; Scan time: ∼4 min

#### Quantitative MRI

An inversion-recovery EPI (IR-EPI) sequence with multiple inversions times (TI) was used to estimate quantitative relaxation time R_1_ (R_1_ = 1/T_1_) in each voxel. The IR-EPI used a slice-shuffling technique to acquire 20 TIs with the first TI = 50 ms and TI interval = 150 ms. A second IR-EPI with reverse-phase encoding direction was also acquired. Other acquisition parameters were voxel size = 2 mm^3^; number of slices = 60; FOV = 20 cm; in-plane/through-plane acceleration = 1/3; Scan time: 1 min and 45 sec. To obtain R_1_ maps, we first performed susceptibility-induced distortion correction on the IR-EPI images using FSL’s top-up^76^ and the IR-EPI acquisition with reverse-phase encoding direction. We then used the distortion corrected images to fit the T_1_ relaxation signal model using a multi-dimensional Levenberg-Marquardt algorithm. In an inversion-recovery sequence, the signal *S(t)* has an exponential decay over time (*t*) with a decay constant *T_1_*:

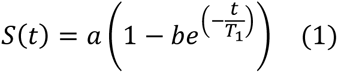

In Eq 1, *a* is a constant that is proportional to the initial magnetization of the voxel and *b* is the effective inversion coefficient of the voxel (for perfect inversion *b* = 2). We applied an absolute value operation on both sides of the equation and used the resulting equation as the fitting model because the magnitude images were used to fit the model. The magnitude images only keep the information about the strength of the signal but not the phase or the sign of the signal. The output of the algorithm is the estimated quantitative T_1_ value per voxel. From the T_1_ estimate, we calculate R_1_ (R_1_ = 1/T_1_) as R_1_ is directly proportional to macromolecular tissue volume.

### Generation of cortical surfaces

We generated gray and white matter tissue segmentations using the T1- and T2-weighted images. Multiple steps were applied to generate an accurate segmentation of each infant’s brain at each timepoint (**Supplementary Fig. 1**): (1) An initial segmentation of gray and white matter was generated from the T1-weighted brain volume using infant FreeSurfer’s automatic segmentation code (infant-recon-all; https://surfer.nmr.mgh.harvard.edu/fswiki/infantFS^45^, (2) a second segmentation was done using both the T1- and T2-weighted anatomical images and the brain extraction toolbox (Brain Extraction and Analysis Toolbox, iBEAT, v-2.0 cloud processing, https://ibeat.wildapricot.org/,^77^ (3) the iBEAT segmentation, which was more accurate than the infant FreeSurfer segmentation, was further manually corrected to fix segmentation errors in the white and gray matter (such as mislabeled white matter voxels or holes) using ITK-SNAP (http://www.itksnap.org/), and (4) the iBEAT corrected segmentation was reinstalled into FreeSurfer and the resulting segmentation in FreeSurfer format was used for all further analyses. A mesh of each infant’s cortical surface was then generated from the boundary of the white and gray matter. This mesh was inflated for visualization of the morphological and macrostructural tissue properties.

### Selecting sulcal folds of interest

To study the morphological development of sulcal folds in the first year of human life, we selected 12 long length sulcal folds (**Fig. 2a-b**): the calcarine (calc), parieto-occipital sulcus (pos), insular sulcus (insula), central sulcus (central), collateral sulcus (CoS), superior temporal sulcus (STS), superior frontal sulcus (SFS), inferior parietal sulcus (IPS), lateral occipital sulcus (LOS), inferior frontal sulcus (IFS), occipital temporal sulcus (OTS), and inferior temporal sulcus (ITS). We chose these sulci as they emerge in different times in utero (16-31 gestational weeks^1^), they are spread across the brain, and they are consistent both across infants and within an infant over time. We used the adult average FreeSurfer brain and cortex-based alignment^46^ to each fold in an individual infant’s brain (**Fig 2b**, example infant, across 4 time points: 0, 3, 6 and 12 months of age).

### Labeling sulci of interest

To delineate the 12 sulci on each infant brain, we first manually labeled each sulcus on the FreeSurfer’s average adult inflated surface (an average surface generated using brain surfaces of 39 adults^46^) using its pial and white matter surfaces to determine sulcal boundaries. We used cortex-based alignment to project each sulcus from the FreeSurfer average cortical surface onto each infant’s cortical surface at each timepoint, using the FreeSurfer function “mri_label2label”. We used the function “mri_label2vol” to convert the projected labels into volume space.

### Quality check on the delineation of sulci

To examine the quality and accuracy of the anatomical placement of the projected sulci from FreeSurfer’s adult average brain onto an individual infant’s brain, we compared the 12 automatically projected sulci to 12 manually delineated sulci in individual infant brains. For this quality check, we randomly selected 10 infants across the four age groups (newborns, 3-month-olds, 6-month-olds, 12-month-olds). Each sulcal fold was hand drawn in the native brain space per infant, per hemisphere. We then measured the overlap between the manually drawn and automatically aligned sulcal labels using dice coefficients. Using a 3-way analysis of variance (ANOVA) with factors: age group (newborns, 3-month-olds, 6-month-olds, 12-month-olds), hemisphere (left/right), and sulcus (*N*=12), and dice coefficients as the dependent measure, we found a significant main effect of sulcus (F_11, 897_ = 106.37, *p* < .0001) and a significant interaction between hemisphere and sulcus (F_11, 897_ = 11.28, *p* < .0001), indicating variability in the accuracy of these projections across different brain sulci. However, there were no significant main effects of age (F_3, 897_ = 0.20, *p* = 0.90), hemisphere (F_1, 897_ = 0.55, *p* = 0.46), or interactions between age and sulcus (F_33, 897_ = 1.13, *p* = 0.28) or age and hemisphere (F_3, 897_ = 1.23, *p* = 0.30). So, while the precision of adult alignment to infants’ cortical surfaces varies with sulcus and hemisphere it does not change with age. This analysis suggests that cortex-based alignment, when applied to infant brains using FreeSurfer’s adult average brain, maintains a consistent level of accuracy across the first year of life, despite significant brain volume changes during this period (**Supplementary Fig. 2**).

### Generation of macrostructural/morphological maps: sulcal depth (SD), cortical thickness (CT), curvature (CU), and sulcal span (SP)

To examine the development of various morphological properties in the 12 sulcal folds, we used FreeSurfer’s automated algorithm to obtain (i) sulcal depth (in mm), (ii) cortical thickness (in mm), (iii) cortical curvature (mm^-1^), and (iv) sulcal span (in mm) for each infant (**Supplementary Fig. 1**). All morphological measurements are generated within each infant’s native brain space and each infant contributed to a single value, per metric, per sulcus.

To obtain the mean depth measures per sulcus and infant, we used the SD maps (lh.sulc/rh.sulc) generated using the FreeSurfer’s auto-segmentation algorithm. SD is measured as the distance between the inflated and white surfaces at each vertex using a movement vectors. For a sulcus, the movement vector points outward (toward the pial surface) and therefore is presented by positive values in FreeSurfer’s lh.sulc and rh.sulc files whereas for a gyral crown, the movement vector points inward and is represented by negative values. Here, we only focused on the sulcal folds. Mean SD (in mm) per sulcus was calculated as the average depth of all voxels in a sulcus. We obtained mean cortical thickness (CT) measured (in mm) as the distance between the gray-white boundary and pial surface along the surface normal. Curvature quantifies 1/r, where r is the radius of an inscribed circle at each vertex (mm^-1^) and determines if the vertex is concave or convex. For sulcal span (SP) is the geodesic distance on the cortical surface from one side of the sulcus to the other side of the sulcus. We estimated the mean SP by calculating the total surface area of each sulcus and dividing it by its length. Surface area is measured as the sum of the areas of all triangles composing the sulcus on the tessellated cortical surface from the area map (lh.area/rh.area); Sulcal length was estimated as distance between the endpoints of the sulcus along its long axis (for instance for the STS – its posterior and anterior points, **Supplementary Fig. 1b**).

### Calculation of mean microstructural tissue density (R_1_) in gray matter

To examine the development of microstructural properties, we used the R_1_ maps per infant and calculated mean R_1_ in the gray matter of each sulcus. Mean R_1_ was calculated as the average R_1_ of all voxels within a sulcus from the gray-white matter boundary to the pial surface. Like morphological parameters, the microstructural properties were also generated from each infant’s native brain space and each infant contributed a single value per sulcus.

### Analysis of development of sulcal depth, sulcal span, thickness, curvature, and R_1_

To quantify developmental effects, we used linear mixed models (LMMs^78^) with the ‘fitlme’ function in MATLAB version 2021b (MathWorks, Inc.). All analyses were done separately for the right and left hemispheres. LMMs allow explicit modeling of both within-subject effects (e.g., longitudinal measurements) and between-subject effects (e.g., cross-sectional data) with unequal number of points per participants, as well as examine main and interactive effects of both continuous (age) and categorical (e.g., sulcus) variables. As developmental effects are larger in the first 6 months than the second 6 months of life, to test development in SD, SP, CT, CU, and R_1_, we fit an LMM relating the parameter of interest to a logarithmic function of infants’ age:

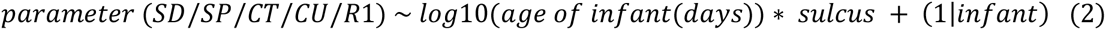

Parameters *SD/SP/CT/CU/R_1_* are the dependent variables, *age of infant* is continuous predictor, *sulcus* is a categorical variable (1 to 12), and the term: *1|infant* indicates a random effect of infant. We ran two types of LMM per parameter: (1) LMM with a random intercept and fixed slope, which allows only the intercepts to vary across areas, and (2) LMM with a random intercept/random slope, which allows both intercepts and slopes to vary across areas and hemispheres. Likelihood tests comparing these two models revealed that the random intercept/random slope model fit the data best (in all model comparisons the latter model was significantly better than the former, likelihood test, *ps* < 0.05). Thus, we report the parameters of LMMs with random intercepts and random slopes. As we found a significant interaction between age and sulcus, we conducted additional LMMs per sulcus to quantify development of each sulcus.

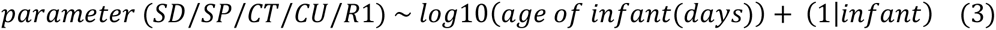

### Prediction of sulcal depth with respect to span, thickness, curvature, and R_1_

To test whether SP, CT, CU, R_1,_ and age predict sulcal depth, we conducted an LMM relating SD to all these parameters. As parameters have different units and ranges, we used normalized values in the models below. Normalization of each parameter (*x*) was as follows: 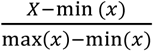, hence the values of model parameters range between 0-1.

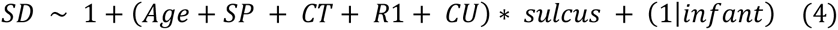

Since we found a global effect of age that was the same across sulci and interactions between sulcus and SP, CT, R1 and CU, we conducted an additional analysis per sulcus to quantify the beta weights of these parameters per sulcus:

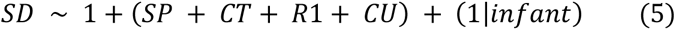

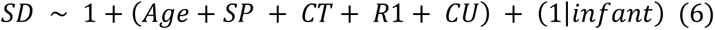

Lastly, we conducted a n-fold-leave-one-out cross-validation approach per sulcus using Eq. 5 to estimate the parameters using data from N-1 infants and use this to predict the mean SD of the left-out infant from their measured SP, CT, CU, and R^1^. Then, we compared the predicted mean depth with actual mean depth values and estimated the root mean squared error (RMSE) per sulcus.

### Generation of sulcal depth, thickness, curvature, and R_1_ profiles

For each sulcus, we generated depth, thickness, curvature, and R_1_ profiles along the long axis of each sulcus. To do this, we first transformed each infant’s SD/CT/CU/R_1_ maps into FreeSurfer’s average cortical space. Next, for a sulcus in the posterior-anterior axis such as the collateral sulcus or the superior temporal sulcus, we divided each sulcus into *N* points based on the number of *z*-coordinates (representing the posterior–anterior axis) along the length of the sulcus. For each *z*-coordinate, we averaged the separately depth, CT, CU, and R_1_ values along the x (lateral– medial) and y (superior–inferior) planes to obtain profiles per micro and macro-level parameter along the sulcus. Specifically, we averaged all the cortical points in a plane orthogonal to the *z*-axis for each *z*-point. Next, to quantitatively test the strength of relationships across sulcal folds and infants and for visualization purpose, we normalized the profiles of SD, CU, R_1_, and CT along the entire length of each sulcus, and calculated the correlations (using Pearson’s correlation coefficient (R)) between SD versus CU/R_1_/CT profiles separately for the left superior temporal sulcus and right collateral sulcus for visualization purpose. To test whether CT, CU, R_1,_ predict sulcal depth along a sulcal fold, we first conducted an LMM relating SD to all these parameters along their anterior-posterior axis:

Since we found a significant interaction between all parameters and sulcus, we conducted an analysis per sulcus to quantify the beta weights of these parameters per sulcus:

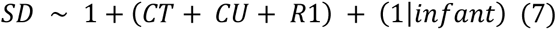

## Code availability

Code to reproduce all the figures in the paper is available on the GitHub repository: https://github.com/VPNL/infants_sulcalmorphology

## Data availability

Source data used in the analyses and to reproduce figures are available on the GitHub repository: https://github.com/VPNL/infants_sulcalmorphology. Requests for raw data should be directed to the corresponding authors, Vaidehi S. Natu and Sarah Tung.

## Acknowledgements

This research was supported by NIH grants R01EY033835, R21 EY030588, Stanford Wu Tsai Neurosciences Institute Big Ideas Grant Phase I and Stanford Wu Tsai Neurosciences Institute Accelerator grants to KGS and by the National Science Foundation Graduate Research Fellowship Program DGE-2146755 to S.S.T. We would also like to thank Kevin S. Weiner and Ethan H. Willbrand for helpful tips for constructing the polar plots.

## Contributions

S.S.T: participant recruitment, data acquisition, data preprocessing, statistical analysis, manuscript writing; X.Y., B.F, & C.T: participant recruitment, data acquisition, data preprocessing; C.M.R, K.D, K.P, A.A, & J.H: data preprocessing; H.W: sequence development; V.S.N & K.G.S: designed, oversaw all components of the study and data analyses, and wrote the manuscript. All co-authors read and approved the submitted manuscript.

---

**Supplementary Figure 1.**
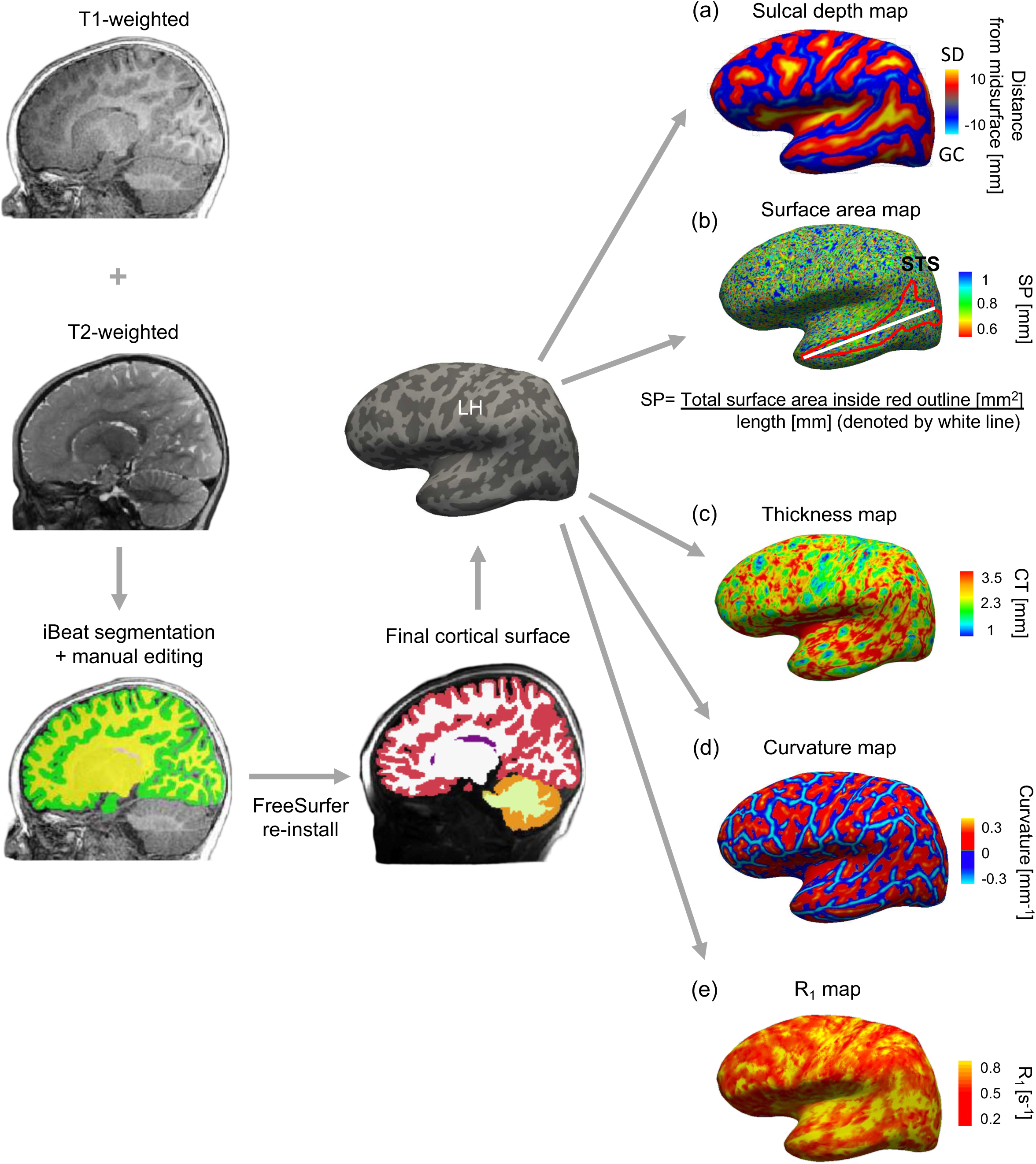
MRI data preprocessing pipeline and microstructural and macrostructural maps. Schematic showing the preprocessing pipeline associated with obtaining the white and gray matter segmentations for generating the cortical surfaces (sample left hemisphere surface) and morphological maps for a sample baby (392 days old) obtained using FreeSurfer’s automated pipeline for (a) sulcal depth [mm], (b) sulcal span [mm] (measured as the total surface area of the STS (for instance) inside the red outline [mm^2^] divided by the length [mm] (denoted by white line) (c), cortical thickness [mm], (d) curvature [mm^-1^], and (e) microstructural quantitative R_1_ [s^-1^] maps aligned to the same anatomical brain volume and cortical surface. All analyses are done for individual baby and timepoint in their native brain space.

**Supplementary Figure 2.**
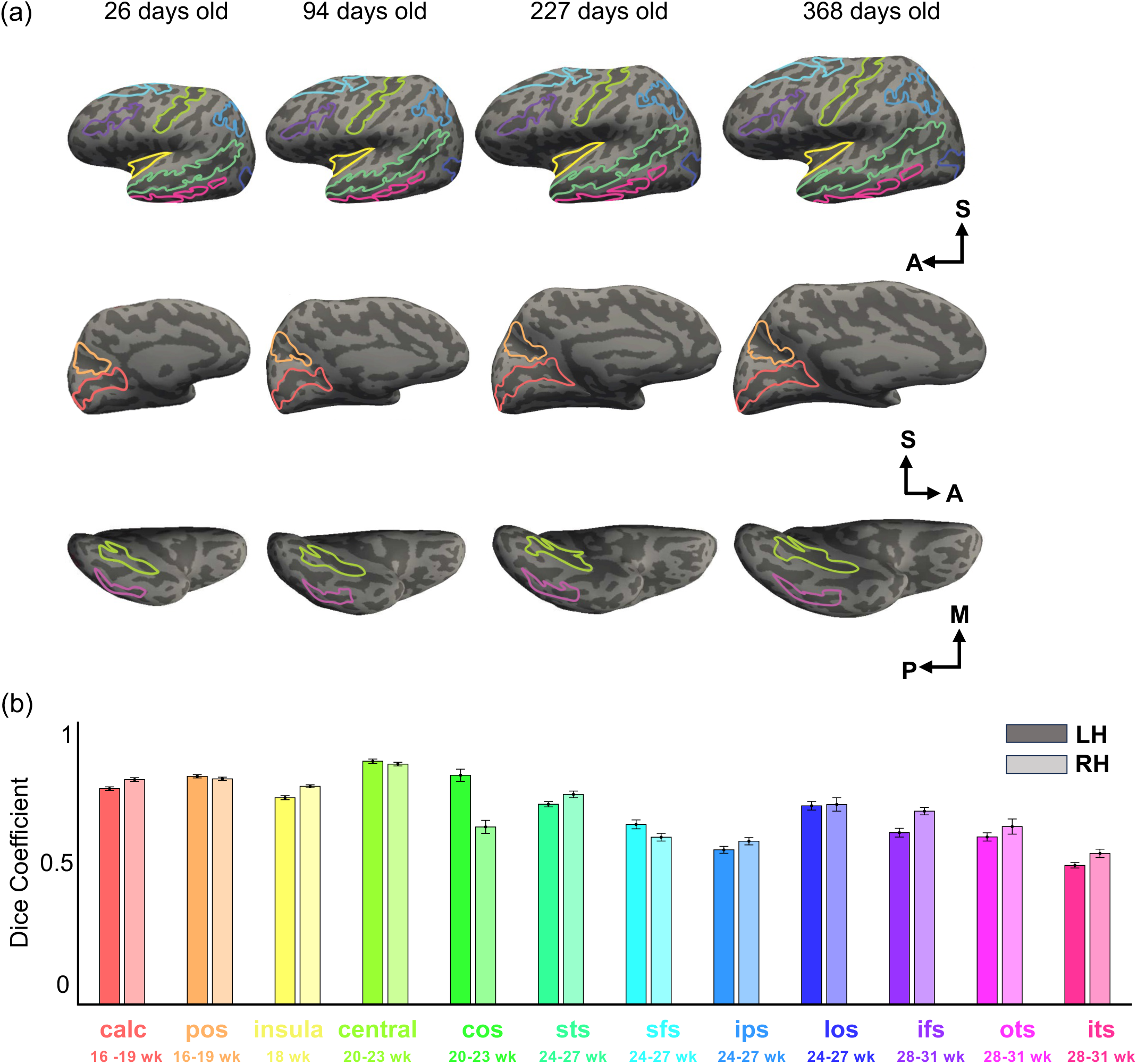
Delineation of sulci. (a) Inflated cortical surfaces of a sample infant’s brain at four postnatal time points (26 days, 94 days, 227 days, and 368 days old) illustrating the progressive development and delineation of sulci across each age within the same infant. Directional arrows indicate the superior (S), anterior (A), medial (M), and posterior (P) orientations. (b) Dice coefficient analysis comparing the accuracy of hand-drawn sulci to projected sulci using FreeSurfer’s adult average template, for both left hemisphere (LH) and right hemisphere (RH), with gestational age ranges indicated beneath each sulcus. Error bars: represent standard error across all participants (*N*=40,10 per age group). *Abbreviations: calc:* calcarine *pos:* parieto-occipital sulcus; *insula:* insular sulcus; *central:* central sulcus; *cos*: collateral sulcus; *sts:* superior temporal sulcus; *sfs:* superior frontal sulcus; *ips*: inferior parietal sulcus; *los:* lateral occipital sulcus, *ifs:* inferior frontal sulcus; *ots:* occipital temporal sulcus; *its:* inferior temporal sulcus. Note: the newborn infant (26 days old) shown here is part of the larger infant data set. However, this timepoint was excluded for the remaining analyses due to unusable qMRI data.

**Supplementary Figure 3.**
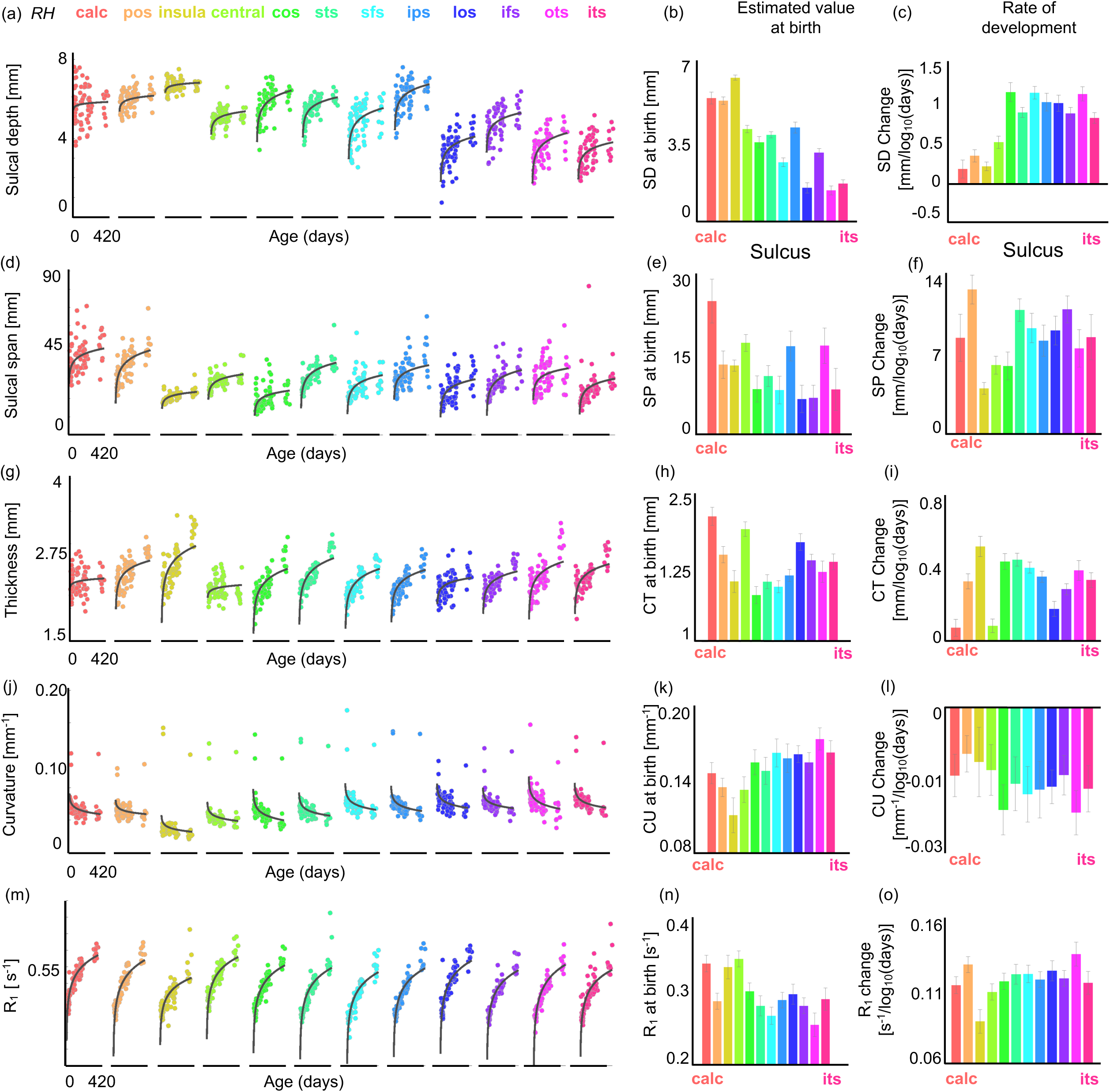
Sulcal folds undergo significant morphological and microstructural changes during the first year of postnatal life in right hemisphere. (a) Sulcal depth (SD) increases logarithmically with age (in days)) in the first year of life in all sulci except for the calcarine and insula. *Each dot*: mean SD per sulcus, per infant. *Line:* Log fit. (b-c) Estimate of SD at birth (intercept) and rate of development of SD (slope) using LMM per sulcus relating mean SD versus log_10_ (participant’s age in days). *Error bars:* standard error on estimates of slopes and intercept. (d-o) Same as in a-c for SP [mm], CT [mm], CU [mm^-1^] and R_1_ [s^-1^]. SP, CT, R_1_ increase with age, but CU decreases, as the folds become less concave. *RH*: Right hemisphere.

**Supplementary Figure 4.**
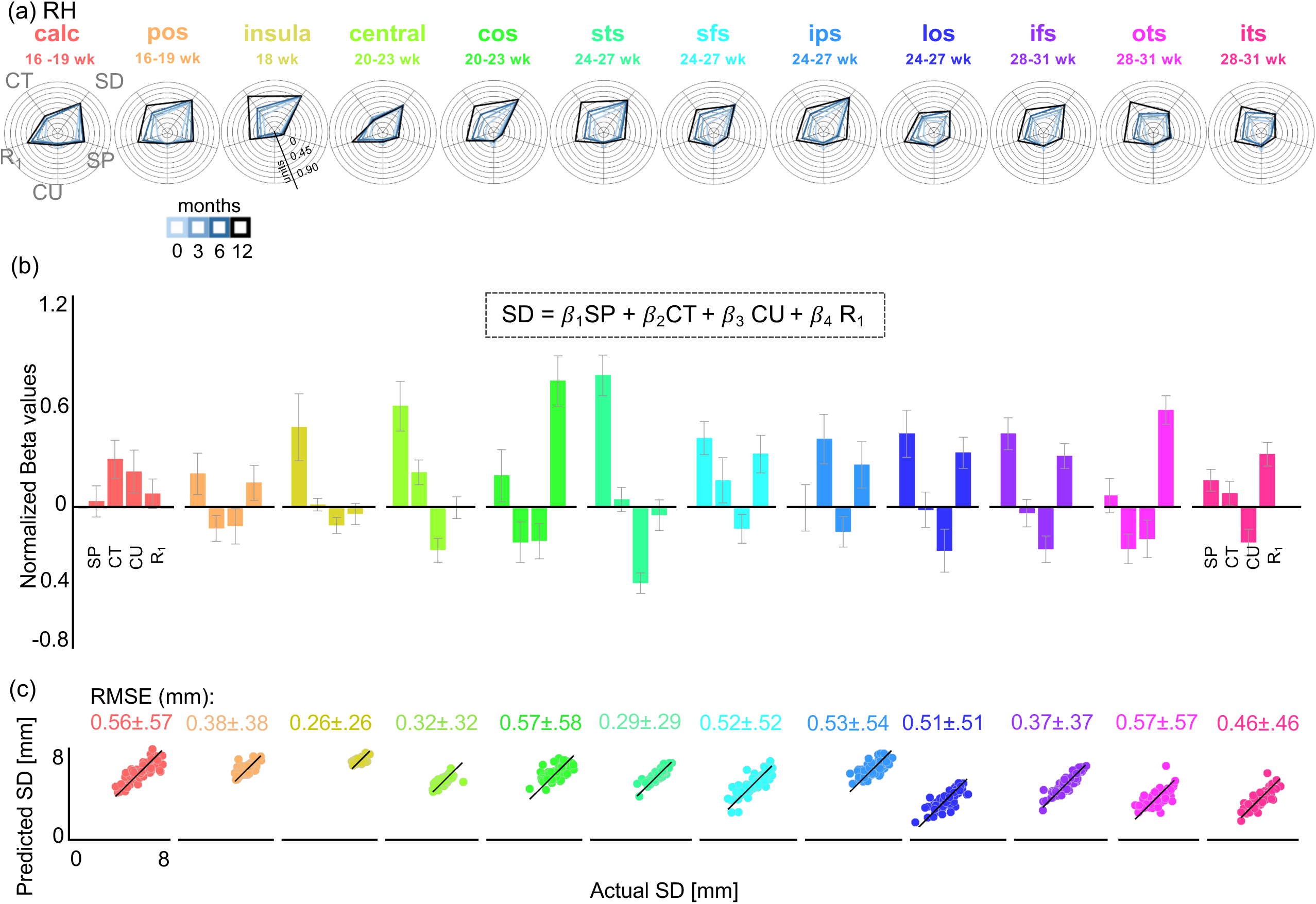
Sulcal depth development can be predicted by a linear combination of SP, CT, CU, and cortical microstructure. (a) Polar plots representing the mean morphological and microstrucural properties across 12 sulci. Four solid lines (light blue – 0 months to dark blue – 12months) represent four age groups. Each concentric circle corresponds to the units shown to the left, which are normalized to allow these metrics to be plotted with respect to each other. (b) Normalized beta values obtained from the ‘best fit’ model (with linear combination of SP, CT, CU, and R_1_) to predict the sulcal development per sulcus. (c) Scatterplots showing predicted depth versus actual depth of each sulcus per participant using the best fit models in (b). RMSE per model is noted above each graph. *RH*: right hemisphere.

**Supplementary Figure 5.**
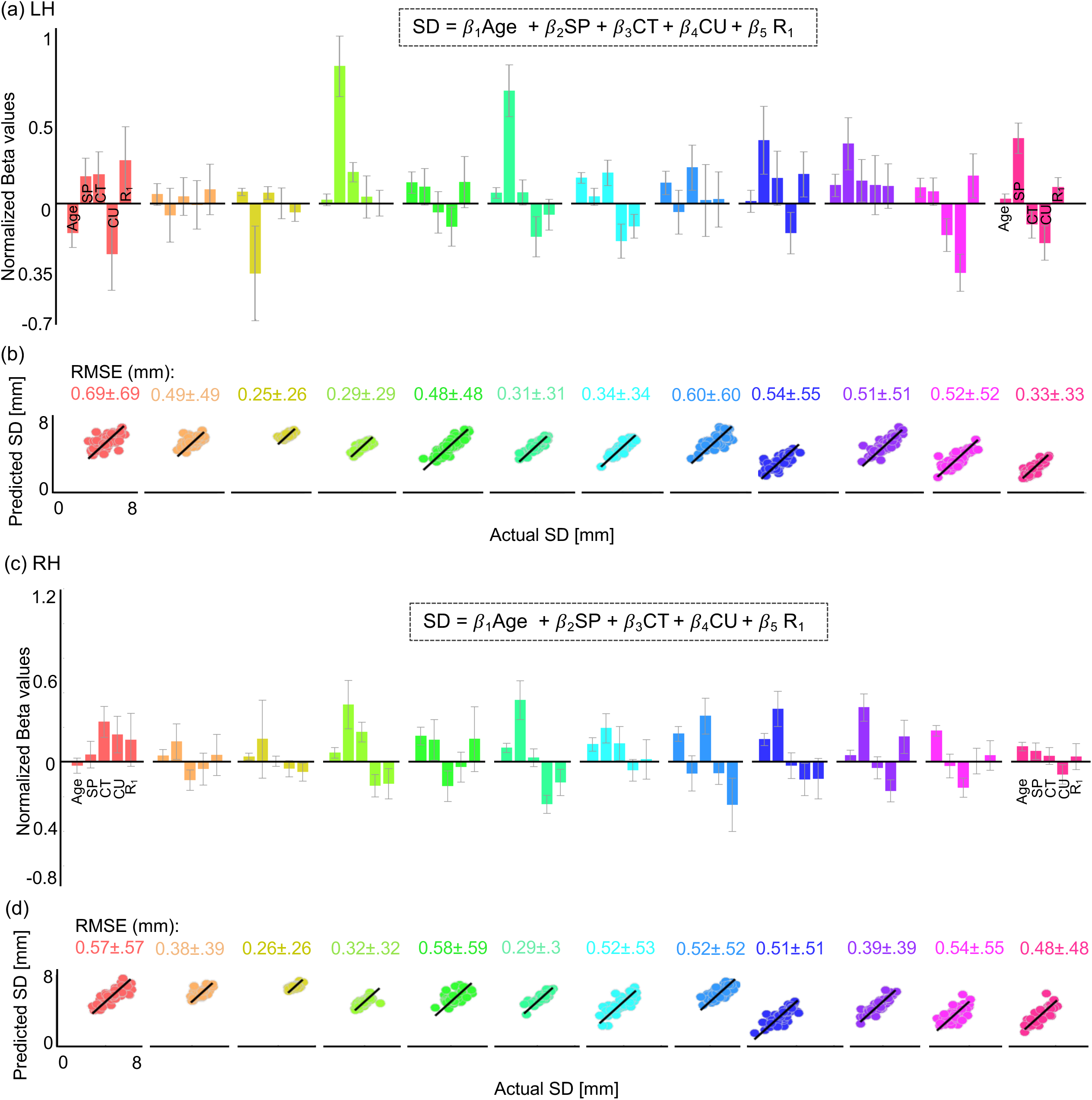
Sulcal depth development can be predicted by a linear combination of Age, SP, CT, CU, and R_1_ for left and right hemispheres. a) Normalized beta values obtained from the ‘best fit’ model (with linear combination of Age, SP, CT, CU, and R_1_) to predict the sulcal development per sulcus in the left hemisphere. (b) Scatterplots showing predicted depth versus actual depth of each sulcus per participant using the best fit models, RMSE per model is noted above each graph. c,d) same as in a,b for the right hemisphere. *LH / RH*: left/right hemisphere.

**Supplementary Figure 6.**
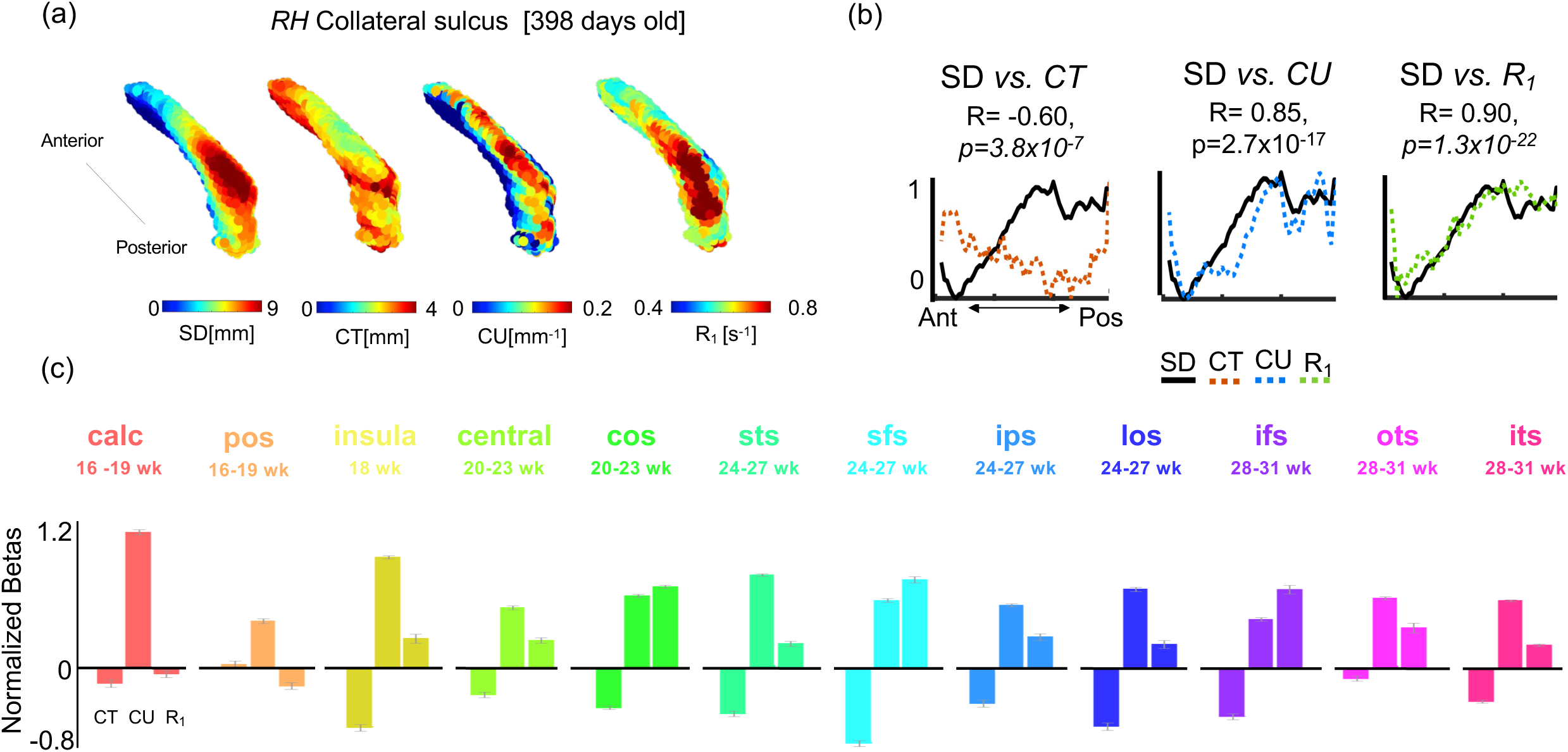
Strong coupling between depth versus micro- and macrostructural properties along the sulcal fold. (a) Three-dimensional maps of right collateral sulcus (CoS) of a sample one year old (398 days) showing corresponding SD, CU, CT, and R_1_ maps (warmer colors represent larger values) (b) Normalized depth (solid black lines), CT (orange dotted lines), CU (blue dotted lines), and R_1_ (green dotted lines) values as a function of distance along the anterior-posterior axis of the CoS showing coupling between depth and each parameter. Correlation (R) and significance values below each subplot relate depth and each parameter. (c) Bar plots showing normalized beta values from a linear model relating SD to CT, CU, and R_1_ along the entire length of the sulcus. *RH*: right hemisphere.

**Supplementary Table 1.**
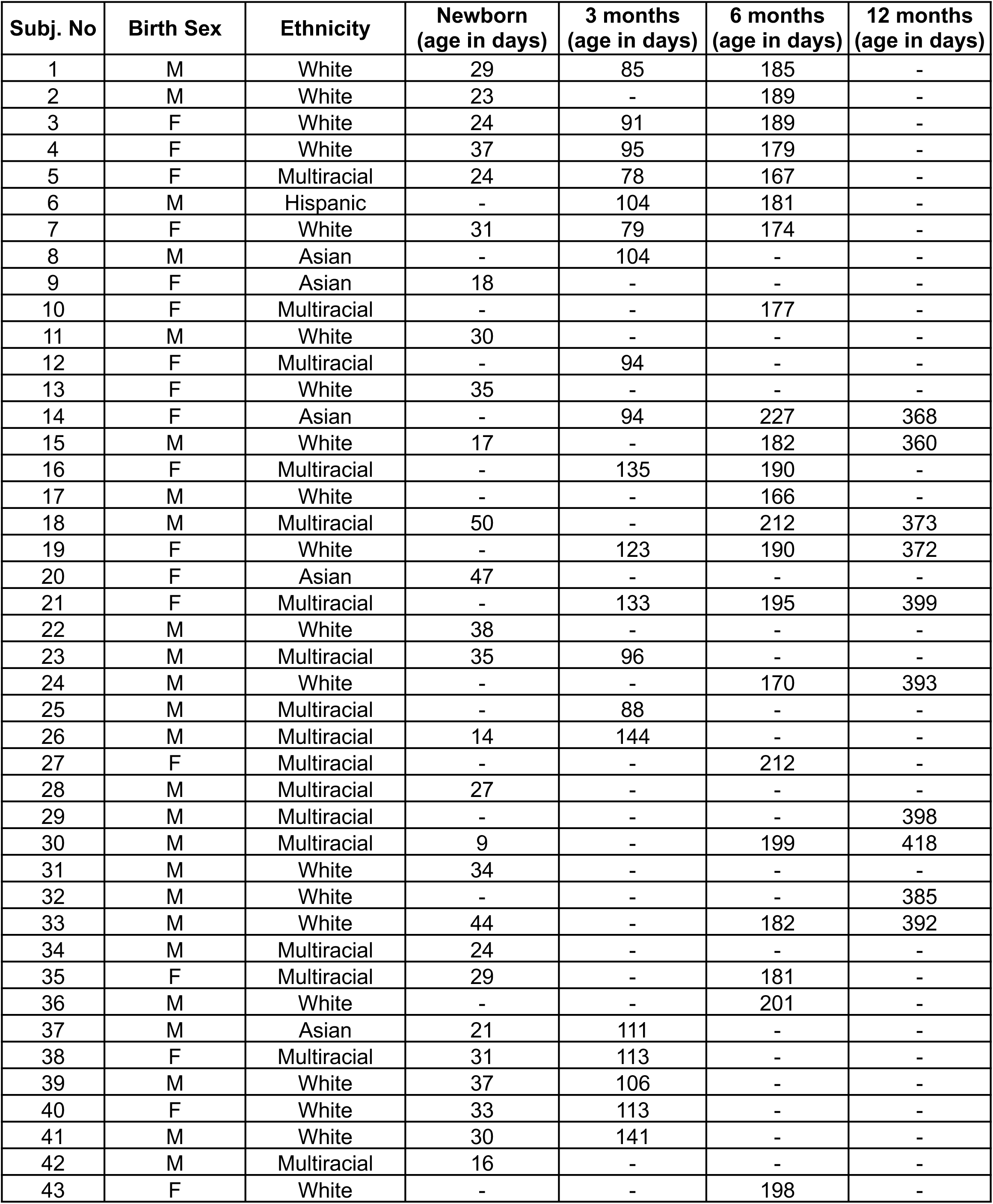
Participant Demographics. Table provides participant demographics, including their birth sex, ethnicity, and their age in days at the time of their MRI sessions at four time points (newborn, 3 months, 6 months, 12 months). Dash (’-’) indicates that a session did not occur or was unusable for the participant at that specific time point.

**Supplementary Table 2.** Statistical significance per linear mixed model (LMM). LMMs quantifying the relationship between parameter and age [log_10_ (age in days)] across the 12 sulci. LMM: *parameter ∼ log10(age of infant) x Sulcus + (1|Infant).* Parameters include: sulcal depth [mm], sulcal span (mm), cortical thickness [mm], curvature [mm^-1^], and microstructure R_1_ [s^-1^]. Three effects are shown per model per hemisphere: main effects of age and sulcus, and interaction between age and sulcus. Statistics: *SE:* Standard error; *DF*: degrees of freedom; *tstats*: t-statistics; *CI* = confidence intervals. *LH/RH:* left/right hemisphere.

| Parameter | Hemi | Effect | $\beta$ Estimate | SE | DF | tStats | pValue | CI lower | CI upper |
| --- | --- | --- | --- | --- | --- | --- | --- | --- | --- |
| SD [mm] | LH | Age | 0.213 | 0.15 | 944 | 1.41 | 0.158 | -0.0828 | 0.508 |
|  | LH | Sulcus | -0.405 | 0.0411 | 944 | -9.87 | 6.38E-22 | -0.486 | -0.325 |
|  | LH | Age x Sulcus | 0.0904 | 0.0204 | 944 | 4.42 | 1.1E-05 | 0.0503 | 0.131 |
|  | RH | Age | 0.194 | 0.147 | 944 | 1.32 | 0.188 | -0.095 | 0.482 |
|  | RH | Sulcus | -0.426 | 0.0402 | 944 | -10.61 | 6.1E-25 | -0.505 | -0.347 |
|  | RH | Age x Sulcus | 0.0916 | 0.02 | 944 | 4.58 | 5.3E-06 | 0.0523 | 0.131 |
| SP [mm] | LH | Age | 8.44 | 1.36 | 944 | 6.21 | 8.1E-10 | 5.77 | 11.1 |
|  | LH | Sulcus | -0.772 | 0.371 | 944 | -2.08 | 0.0377 | -1.5 | -0.044 |
|  | LH | Age x Sulcus | -0.00943 | 0.185 | 944 | -0.05 | 0.959 | -0.372 | 0.353 |
|  | RH | Age | 7.97 | 1.37 | 944 | 5.82 | 8.3E-09 | 5.28 | 10.7 |
|  | RH | Sulcus | -0.857 | 0.374 | 944 | -2.29 | 0.0222 | -1.59 | -0.123 |
|  | RH | Age x Sulcus | 0.108 | 0.186 | 944 | 0.58 | 0.562 | -0.258 | 0.474 |
| CT [mm] | LH | Age | 0.255 | 0.0311 | 944 | 8.19 | 8.3E-16 | 0.194 | 0.315 |
|  | LH | Sulcus | -0.0221 | 0.00809 | 944 | -2.73 | 0.00652 | -0.0379 | -0.00618 |
|  | LH | Age x Sulcus | 0.00959 | 0.00403 | 944 | 2.38 | 0.0175 | 0.00169 | 0.0175 |
|  | RH | Age | 0.259 | 0.0316 | 944 | 8.19 | 8.7E-16 | 0.197 | 0.321 |
|  | RH | Sulcus | -0.0234 | 0.00863 | 944 | -2.71 | 0.00686 | -0.0403 | -0.00645 |
|  | RH | Age x Sulcus | 0.00921 | 0.0043 | 944 | 2.14 | 0.0323 | 0.000779 | 0.0176 |
| CU [mm<sup>-1</sup>] | LH | Age | -0.00956 | 0.00198 | 944 | -4.82 | 1.6E-06 | -0.0135 | -0.00567 |
|  | LH | Sulcus | 0.00354 | 0.000541 | 944 | 6.53 | 1E-10 | 0.00247 | 0.0046 |
|  | LH | Age x Sulcus | -0.000673 | 0.000269 | 944 | -2.5 | 0.0127 | -0.0012 | -0.000144 |
|  | RH | Age | -0.00664 | 0.00293 | 944 | -2.27 | 0.0236 | -0.0124 | -0.000892 |
|  | RH | Sulcus | 0.00363 | 0.00074 | 944 | 4.9 | 1.1E-06 | 0.00218 | 0.00508 |
|  | RH | Age x Sulcus | -0.000612 | 0.000368 | 944 | -1.66 | 0.0966 | -0.00134 | 0.00011 |
| R<sub>1</sub> [s<sup>-1</sup>] | LH | Age | 0.104 | 0.0039 | 944 | 26.59 | 9.73E-117 | 0.0962 | 0.111 |
|  | LH | Sulcus | -0.00586 | 0.000992 | 944 | -5.91 | 4.79E-09 | -0.00781 | -0.00391 |
|  | LH | Age x Sulcus | 0.00154 | 0.000494 | 944 | 3.11 | 0.0019 | 0.000568 | 0.00251 |
|  | RH | Age | 0.106 | 0.00442 | 944 | 23.92 | 3E-99 | 0.0971 | 0.115 |
|  | RH | Sulcus | -0.0054 | 0.00112 | 944 | -4.8 | 1.8E-06 | -0.0076 | -0.00319 |
|  | RH | Age x Sulcus | 0.00138 | 0.000559 | 944 | 2.47 | 0.0138 | 0.000282 | 0.00248 |

**Supplementary Table 3.** Statistical significance of linear mixed models (LMMs), quantifying the relationship between mean sulcal depth [mm] and age [log_10_ (age in days)] for 12 sulci: *SD ∼ log10(age) + (1|Infan*t). Related to **Figs. 2d-f and Supplementary Figs. 3a-c**. Intercept units: millimeters [mm]; Slope units: [mm/log_10_ (age in days)]. Statistics: *Rsquared* = proportion of variance explained; *DF*: degrees of freedom; *tstats*: t-statistics; *CI* = confidence intervals. *LH/RH:* left/right hemisphere.

| Sulcus | Hemisphere | Intercept | Slope | pValue | Rsquared | DF | tstats | CI Lower | CI Upper |
| --- | --- | --- | --- | --- | --- | --- | --- | --- | --- |
| calcarine | LH | 5.93 | -0.15 | 0.31 | 0.8 | 77 | -1.02 | -0.436 | 0.14 |
| pos | LH | 4.72 | 0.5 | 2.4E-05 | 0.75 | 77 | 4.5 | 0.276 | 0.715 |
| insula | LH | 5.78 | 0.26 | 1.1E-06 | 0.84 | 77 | 5.3 | 0.165 | 0.364 |
| central | LH | 3.96 | 0.59 | 4.9E-10 | 0.73 | 77 | 7.12 | 0.422 | 0.749 |
| cos | LH | 2.96 | 1.16 | 3.1E-18 | 0.94 | 77 | 11.42 | 0.96 | 1.37 |
| sts | LH | 3.24 | 1.01 | 8.4E-26 | 0.94 | 77 | 15.76 | 0.882 | 1.14 |
| sfs | LH | 2.76 | 1.04 | 2.4E-28 | 0.97 | 77 | 17.36 | 0.923 | 1.16 |
| ips | LH | 3.97 | 0.94 | 5.4E-12 | 0.89 | 77 | 8.14 | 0.708 | 1.17 |
| loc | LH | 1.45 | 1.00 | 6E-11 | 0.8 | 77 | 7.6 | 0.737 | 1.26 |
| ifs | LH | 2.89 | 1.19 | 1.2E-18 | 0.93 | 77 | 11.64 | 0.989 | 1.4 |
| ots | LH | 1.42 | 1.13 | 9.6E-18 | 0.93 | 77 | 11.15 | 0.93 | 1.33 |
| its | LH | 1.21 | 0.75 | 6E-15 | 0.91 | 77 | 9.68 | 0.594 | 0.901 |
| calcarine | RH | 5.36 | 0.19 | 0.105 | 0.92 | 77 | 1.64 | -0.0396 | 0.412 |
| pos | RH | 5.25 | 0.35 | 1.1E-05 | 0.85 | 77 | 4.7 | 0.202 | 0.499 |
| insula | RH | 6.25 | 0.22 | 8.3E-05 | 0.76 | 77 | 4.16 | 0.115 | 0.325 |
| central | RH | 4.02 | 0.52 | 2.2E-09 | 0.77 | 77 | 6.77 | 0.366 | 0.671 |
| cos | RH | 3.45 | 1.14 | 6.2E-15 | 0.83 | 77 | 9.67 | 0.905 | 1.37 |
| sts | RH | 3.76 | 0.89 | 3.2E-23 | 0.94 | 77 | 14.21 | 0.763 | 1.01 |
| sfs | RH | 2.57 | 1.13 | 5.6E-24 | 0.97 | 77 | 14.65 | 0.979 | 1.29 |
| ips | RH | 4.08 | 1.02 | 4.7E-14 | 0.87 | 77 | 9.21 | 0.797 | 1.24 |
| loc | RH | 1.46 | 1.00 | 1.6E-15 | 0.92 | 77 | 9.98 | 0.803 | 1.2 |
| ifs | RH | 3.00 | 0.88 | 2.8E-19 | 0.96 | 77 | 11.98 | 0.731 | 1.02 |
| ots | RH | 1.35 | 1.12 | 1.4E-20 | 0.93 | 77 | 12.7 | 0.942 | 1.29 |
| its | RH | 1.65 | 0.82 | 8E-20 | 0.97 | 77 | 12.28 | 0.687 | 0.953 |

**Supplementary Table 4.** Statistical significance of linear mixed models (LMMs) quantifying the relationship between mean sulcal span [mm] and age [log_10_ (age in days)] for 12 sulci: *SP ∼ log10(age) + (1|Infant)*. Related to **Figs. 3a-c and Supplementary Figs. 3d-f**. Intercept units: millimeters [mm]; Slope units: [mm/log_10_ (age in days)]. Statistics: *Rsquared* = proportion of variance explained; *DF*: degrees of freedom; *tstats*: t-statistics; *CI* = confidence intervals. *LH/RH:* left/right hemisphere.

| Sulcus | Hemisphere | Intercept | Slope | pValue | Rsquared | DF | tstats | CI Lower | CI Upper |
| --- | --- | --- | --- | --- | --- | --- | --- | --- | --- |
| calcarine | LH | 23.8 | 7.84 | 0.00015 | 0.61 | 77 | 3.99 | 3.93 | 11.8 |
| pos | LH | 15.8 | 10.3 | 3.4E-17 | 0.91 | 77 | 10.86 | 8.41 | 12.2 |
| insula | LH | 11.5 | 4.3 | 2.1E-25 | 0.78 | 77 | 15.52 | 3.75 | 4.85 |
| central | LH | 15.8 | 6.52 | 3.8E-20 | 0.87 | 77 | 12.46 | 5.48 | 7.56 |
| cos | LH | 12.5 | 6.08 | 0.00071 | 0.25 | 77 | 3.53 | 2.65 | 9.52 |
| sts | LH | 12.7 | 10.72 | 1.4E-27 | 0.94 | 77 | 16.88 | 9.45 | 12 |
| sfs | LH | 11.1 | 10.32 | 1.6E-07 | 0.47 | 77 | 5.76 | 6.75 | 13.9 |
| ips | LH | 9.79 | 11.61 | 1.9E-12 | 0.82 | 77 | 8.38 | 8.85 | 14.4 |
| loc | LH | 11.1 | 6.13 | 2E-10 | 0.83 | 77 | 7.33 | 4.46 | 7.79 |
| ifs | LH | 6.23 | 9.47 | 2.5E-13 | 0.76 | 77 | 8.83 | 7.33 | 11.6 |
| ots | LH | 16.5 | 8.47 | 5.3E-05 | 0.48 | 77 | 4.28 | 4.53 | 12.4 |
| its | LH | 3.54 | 7.77 | 3E-09 | 0.55 | 77 | 6.71 | 5.47 | 10.1 |
| calcarine | RH | 25.3 | 8.48 | 9.4E-05 | 0.35 | 77 | 4.12 | 4.38 | 12.6 |
| pos | RH | 13.2 | 12.7 | 1.1E-15 | 0.81 | 77 | 10.07 | 10.2 | 15.2 |
| insula | RH | 13.1 | 4.05 | 2.8E-11 | 0.52 | 77 | 7.78 | 3.01 | 5.09 |
| central | RH | 17.4 | 6.09 | 2.5E-11 | 0.63 | 77 | 7.8 | 4.54 | 7.65 |
| cos | RH | 8.62 | 6.02 | 4E-06 | 0.63 | 77 | 4.97 | 3.6 | 8.43 |
| sts | RH | 11.1 | 10.91 | 8.9E-18 | 0.71 | 77 | 11.17 | 8.96 | 12.9 |
| sfs | RH | 8.42 | 9.34 | 3.5E-10 | 0.55 | 77 | 7.2 | 6.75 | 11.9 |
| ips | RH | 16.8 | 8.23 | 4.9E-08 | 0.79 | 77 | 6.05 | 5.52 | 10.9 |
| loc | RH | 6.74 | 9.11 | 3.4E-10 | 0.76 | 77 | 7.2 | 6.59 | 11.6 |
| ifs | RH | 6.95 | 10.97 | 3.3E-14 | 0.76 | 77 | 9.29 | 8.62 | 13.3 |
| ots | RH | 16.8 | 7.56 | 1.4E-05 | 0.67 | 77 | 4.64 | 4.32 | 10.8 |
| its | RH | 8.52 | 8.55 | 4.5E-05 | 0.23 | 77 | 4.33 | 4.61 | 12.5 |

**Supplementary Table 5.** Statistical significance of linear mixed models (LMMs) quantifying the relationship between mean cortical thickness [mm] and age [log_10_ (age in days)] for 12 sulci: *CT ∼ log10(age) + (1|infant).* Related to **Figs. 3d-f and Supplementary Figs. 3g-i**. Intercept units: millimeters [mm]; Slope units: [mm/log_10_ (age in days)]. Statistics: *Rsquared* = proportion of variance explained; *DF*: degrees of freedom; *tstats*: t-statistics; *CI* = confidence intervals. *LH/RH:* left/right hemisphere.

| Sulcus | Hemisphere | Intercept | Slope | P value | Rsquared | DF | tstats | CI Lower | CI Upper |
| --- | --- | --- | --- | --- | --- | --- | --- | --- | --- |
| calcarine | LH | 2.33 | 0.03 | 0.53 | 0.04 | 77 | 0.63 | -0.0565 | 0.109 |
| pos | LH | 1.89 | 0.3 | 4E-10 | 0.43 | 77 | 7.17 | 0.218 | 0.386 |
| insula | LH | 1.56 | 0.53 | 1.6E-16 | 0.58 | 77 | 10.51 | 0.428 | 0.628 |
| central | LH | 2.22 | 0.05 | 0.232 | 0.09 | 77 | 1.2 | -0.034 | 0.138 |
| cos | LH | 1.36 | 0.48 | 1.2E-18 | 0.79 | 77 | 11.65 | 0.396 | 0.559 |
| sts | LH | 1.72 | 0.37 | 2.1E-14 | 0.54 | 77 | 9.39 | 0.292 | 0.45 |
| sfs | LH | 1.58 | 0.41 | 2.9E-21 | 0.74 | 77 | 13.09 | 0.35 | 0.476 |
| ips | LH | 1.62 | 0.4 | 4E-16 | 0.7 | 77 | 10.29 | 0.321 | 0.475 |
| loc | LH | 1.88 | 0.23 | 1.7E-10 | 0.66 | 77 | 7.36 | 0.169 | 0.294 |
| ifs | LH | 1.68 | 0.35 | 1.1E-15 | 0.65 | 77 | 10.05 | 0.278 | 0.416 |
| ots | LH | 1.71 | 0.35 | 1.6E-09 | 0.51 | 77 | 6.85 | 0.245 | 0.446 |
| its | LH | 2.01 | 0.25 | 5.6E-08 | 0.32 | 77 | 6.02 | 0.169 | 0.336 |
| calcarine | RH | 2.27 | 0.08 | 0.101 | 0.16 | 77 | 1.66 | -0.0151 | 0.165 |
| pos | RH | 1.88 | 0.33 | 1E-11 | 0.63 | 77 | 8.00 | 0.246 | 0.409 |
| insula | RH | 1.61 | 0.52 | 2.1E-14 | 0.61 | 77 | 9.39 | 0.407 | 0.627 |
| central | RH | 2.14 | 0.09 | 0.0222 | 0.16 | 77 | 2.33 | 0.0126 | 0.158 |
| cos | RH | 1.47 | 0.44 | 1.3E-15 | 0.65 | 77 | 10.02 | 0.349 | 0.522 |
| sts | RH | 1.61 | 0.45 | 2.3E-20 | 0.66 | 77 | 12.59 | 0.376 | 0.517 |
| sfs | RH | 1.56 | 0.4 | 5.7E-22 | 0.78 | 77 | 13.49 | 0.343 | 0.462 |
| ips | RH | 1.67 | 0.35 | 2.1E-18 | 0.74 | 77 | 11.51 | 0.292 | 0.414 |
| loc | RH | 2.01 | 0.18 | 5.2E-05 | 0.45 | 77 | 4.28 | 0.0951 | 0.26 |
| ifs | RH | 1.83 | 0.29 | 1.9E-14 | 0.59 | 77 | 9.41 | 0.225 | 0.346 |
| ots | RH | 1.71 | 0.39 | 3.2E-10 | 0.48 | 77 | 7.22 | 0.281 | 0.495 |
| its | RH | 1.81 | 0.34 | 2E-12 | 0.46 | 77 | 8.37 | 0.256 | 0.415 |

**Supplementary Table 6.**
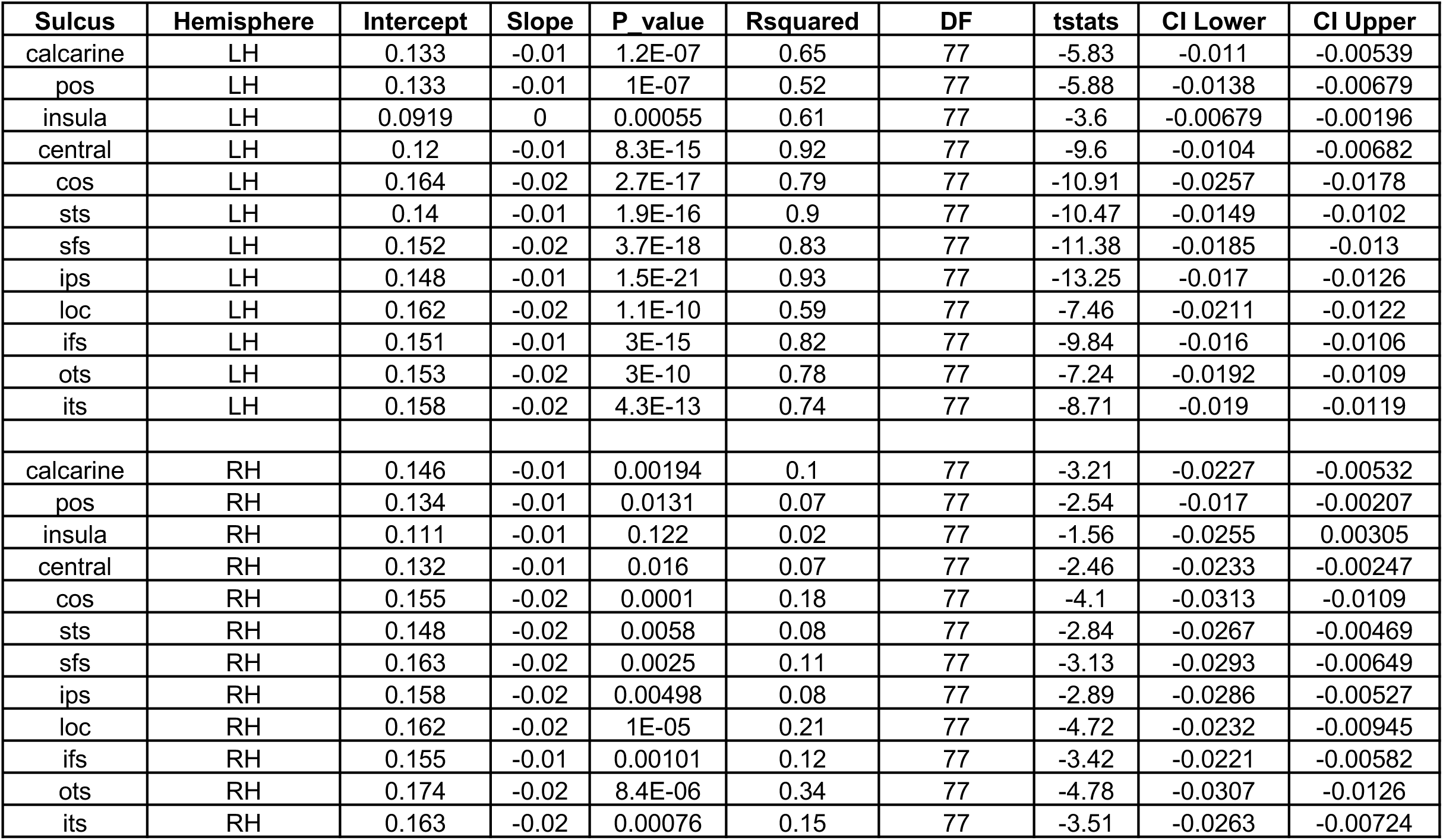
Statistical significance of linear mixed models (LMMs) quantifying the relationship between mean curvature [mm^-1^] and age [log_10_ (age in days)] for 12 sulci: *CU ∼ log10(age) + (1|infant).* Related to **Figs. 3g-i and Supplementary Figs. 3j-l**. Intercept units: millimeters [mm^-1^]; Slope units: [mm^-1^ /log_10_ (age in days)]. Statistics: *Rsquared* = proportion of variance explained; *DF*: degrees of freedom; *tstats*: t-statistics; *CI* = confidence intervals. *LH/RH:* left/right hemisphere.

| Sulcus | Hemisphere | Intercept | Slope | P value | Rsquared | DF | tstats | CI Lower | CI Upper |
| --- | --- | --- | --- | --- | --- | --- | --- | --- | --- |
| calcarine | LH | 0.133 | -0.01 | 1.2E-07 | 0.65 | 77 | -5.83 | -0.011 | -0.00539 |
| pos | LH | 0.133 | -0.01 | 1E-07 | 0.52 | 77 | -5.88 | -0.0138 | -0.00679 |
| insula | LH | 0.0919 | 0 | 0.00055 | 0.61 | 77 | -3.6 | -0.00679 | -0.00196 |
| central | LH | 0.12 | -0.01 | 8.3E-15 | 0.92 | 77 | -9.6 | -0.0104 | -0.00682 |
| cos | LH | 0.164 | -0.02 | 2.7E-17 | 0.79 | 77 | -10.91 | -0.0257 | -0.0178 |
| sts | LH | 0.14 | -0.01 | 1.9E-16 | 0.9 | 77 | -10.47 | -0.0149 | -0.0102 |
| sfs | LH | 0.152 | -0.02 | 3.7E-18 | 0.83 | 77 | -11.38 | -0.0185 | -0.013 |
| ips | LH | 0.148 | -0.01 | 1.5E-21 | 0.93 | 77 | -13.25 | -0.017 | -0.0126 |
| loc | LH | 0.162 | -0.02 | 1.1E-10 | 0.59 | 77 | -7.46 | -0.0211 | -0.0122 |
| ifs | LH | 0.151 | -0.01 | 3E-15 | 0.82 | 77 | -9.84 | -0.016 | -0.0106 |
| ots | LH | 0.153 | -0.02 | 3E-10 | 0.78 | 77 | -7.24 | -0.0192 | -0.0109 |
| its | LH | 0.158 | -0.02 | 4.3E-13 | 0.74 | 77 | -8.71 | -0.019 | -0.0119 |
| calcarine | RH | 0.146 | -0.01 | 0.00194 | 0.1 | 77 | -3.21 | -0.0227 | -0.00532 |
| pos | RH | 0.134 | -0.01 | 0.0131 | 0.07 | 77 | -2.54 | -0.017 | -0.00207 |
| insula | RH | 0.111 | -0.01 | 0.122 | 0.02 | 77 | -1.56 | -0.0255 | 0.00305 |
| central | RH | 0.132 | -0.01 | 0.016 | 0.07 | 77 | -2.46 | -0.0233 | -0.00247 |
| cos | RH | 0.155 | -0.02 | 0.0001 | 0.18 | 77 | -4.1 | -0.0313 | -0.0109 |
| sts | RH | 0.148 | -0.02 | 0.0058 | 0.08 | 77 | -2.84 | -0.0267 | -0.00469 |
| sfs | RH | 0.163 | -0.02 | 0.0025 | 0.11 | 77 | -3.13 | -0.0293 | -0.00649 |
| ips | RH | 0.158 | -0.02 | 0.00498 | 0.08 | 77 | -2.89 | -0.0286 | -0.00527 |
| loc | RH | 0.162 | -0.02 | 1E-05 | 0.21 | 77 | -4.72 | -0.0232 | -0.00945 |
| ifs | RH | 0.155 | -0.01 | 0.00101 | 0.12 | 77 | -3.42 | -0.0221 | -0.00582 |
| ots | RH | 0.174 | -0.02 | 8.4E-06 | 0.34 | 77 | -4.78 | -0.0307 | -0.0126 |
| its | RH | 0.163 | -0.02 | 0.00076 | 0.15 | 77 | -3.51 | -0.0263 | -0.00724 |

**Supplementary Table 7.**
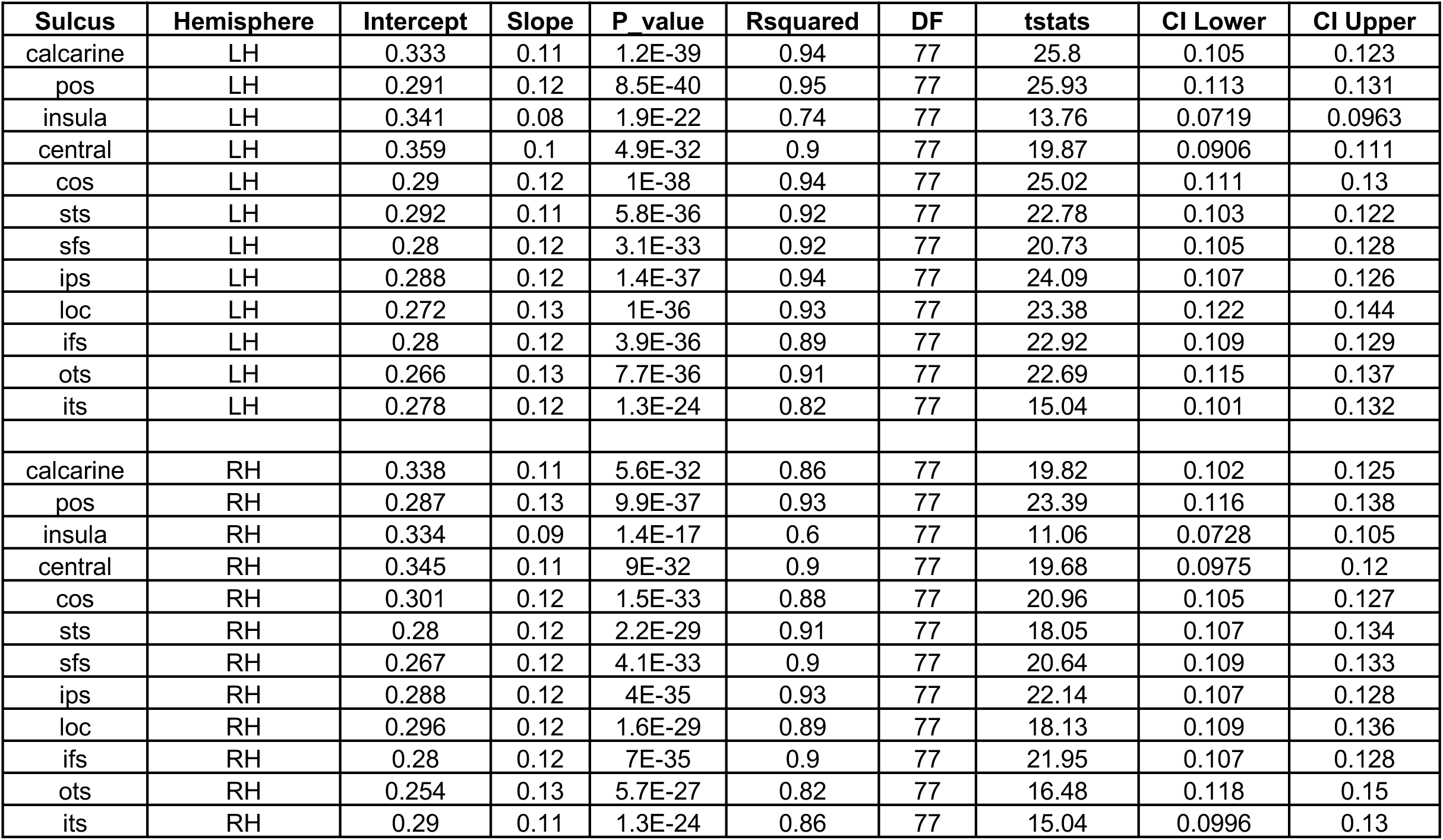
Statistical significance of linear mixed models (LMMs) quantifying the relationship between mean cortical microstructure [s^-1^] and age [log_10_ (age in days)] for 12 sulci: *R_1_ ∼ log10(age) +.(1|infant).* Related to **Figs. 3j-l and Supplementary Figs. 3m-o**. Intercept units: millimeters [s^-1^]; Slope units: [s^-1^ /log_10_ (age in days)]. Statistics: *Rsquared* = proportion of variance explained; *DF*: degrees of freedom; *tstats*: t-statistics; *CI* = confidence intervals. *LH/RH:* left/right hemisphere.

| Sulcus | Hemisphere | Intercept | Slope | P value | Rsquared | DF | tstats | CI Lower | CI Upper |
| --- | --- | --- | --- | --- | --- | --- | --- | --- | --- |
| calcarine | LH | 0.333 | 0.11 | 1.2E-39 | 0.94 | 77 | 25.8 | 0.105 | 0.123 |
| pos | LH | 0.291 | 0.12 | 8.5E-40 | 0.95 | 77 | 25.93 | 0.113 | 0.131 |
| insula | LH | 0.341 | 0.08 | 1.9E-22 | 0.74 | 77 | 13.76 | 0.0719 | 0.0963 |
| central | LH | 0.359 | 0.1 | 4.9E-32 | 0.9 | 77 | 19.87 | 0.0906 | 0.111 |
| cos | LH | 0.29 | 0.12 | 1E-38 | 0.94 | 77 | 25.02 | 0.111 | 0.13 |
| sts | LH | 0.292 | 0.11 | 5.8E-36 | 0.92 | 77 | 22.78 | 0.103 | 0.122 |
| sfs | LH | 0.28 | 0.12 | 3.1E-33 | 0.92 | 77 | 20.73 | 0.105 | 0.128 |
| ips | LH | 0.288 | 0.12 | 1.4E-37 | 0.94 | 77 | 24.09 | 0.107 | 0.126 |
| loc | LH | 0.272 | 0.13 | 1E-36 | 0.93 | 77 | 23.38 | 0.122 | 0.144 |
| ifs | LH | 0.28 | 0.12 | 3.9E-36 | 0.89 | 77 | 22.92 | 0.109 | 0.129 |
| ots | LH | 0.266 | 0.13 | 7.7E-36 | 0.91 | 77 | 22.69 | 0.115 | 0.137 |
| its | LH | 0.278 | 0.12 | 1.3E-24 | 0.82 | 77 | 15.04 | 0.101 | 0.132 |
| calcarine | RH | 0.338 | 0.11 | 5.6E-32 | 0.86 | 77 | 19.82 | 0.102 | 0.125 |
| pos | RH | 0.287 | 0.13 | 9.9E-37 | 0.93 | 77 | 23.39 | 0.116 | 0.138 |
| insula | RH | 0.334 | 0.09 | 1.4E-17 | 0.6 | 77 | 11.06 | 0.0728 | 0.105 |
| central | RH | 0.345 | 0.11 | 9E-32 | 0.9 | 77 | 19.68 | 0.0975 | 0.12 |
| cos | RH | 0.301 | 0.12 | 1.5E-33 | 0.88 | 77 | 20.96 | 0.105 | 0.127 |
| sts | RH | 0.28 | 0.12 | 2.2E-29 | 0.91 | 77 | 18.05 | 0.107 | 0.134 |
| sfs | RH | 0.267 | 0.12 | 4.1E-33 | 0.9 | 77 | 20.64 | 0.109 | 0.133 |
| ips | RH | 0.288 | 0.12 | 4E-35 | 0.93 | 77 | 22.14 | 0.107 | 0.128 |
| loc | RH | 0.296 | 0.12 | 1.6E-29 | 0.89 | 77 | 18.13 | 0.109 | 0.136 |
| ifs | RH | 0.28 | 0.12 | 7E-35 | 0.9 | 77 | 21.95 | 0.107 | 0.128 |
| ots | RH | 0.254 | 0.13 | 5.7E-27 | 0.82 | 77 | 16.48 | 0.118 | 0.15 |
| its | RH | 0.29 | 0.11 | 1.3E-24 | 0.86 | 77 | 15.04 | 0.0996 | 0.13 |

**Supplementary Table 8.**
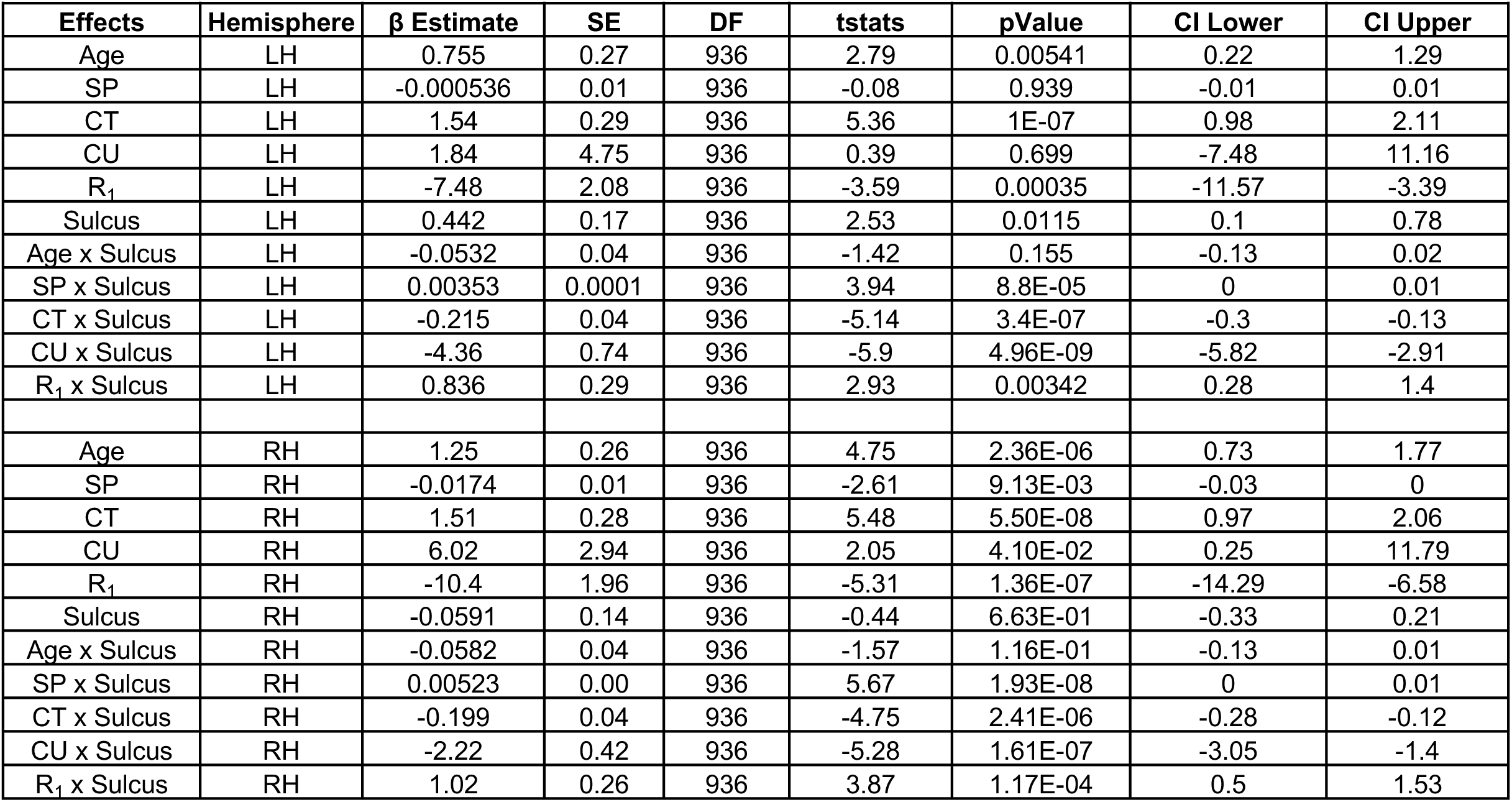
Statistical significance and estimates of the linear mixed model (LMMs): *SD ∼ 1 + (Age + SP + CT + CU + R_1_) x Sulcus + (1|Infant).* Statistics: *β:* beta coefficients; *SE:* standard errors; *DF*: degrees of freedom; *tstats*: t-statistics; *CI:* confidence intervals. *LH/RH:* left/right hemisphere.

**Supplementary Table 9.** Normalized beta coefficients *(β)* and standard errors *(SE)* for sulcal span (SP), cortical thickness (CT), curvature, (CU), and cortical microstructure (R_1_) across sulci in the left/right hemispheres (LH/RH). The beta coefficients represent the relationship between sulcal depth and each mean parameter as described by the model: *SD = 1 + β_1_SP + β_2_CT + β_3_ CU + β_4_R_1_ + (1|Infant)*.

| Sulcus | Hemi | $\beta$ SP | SE SP | $\beta$ CT | SE CT | $\beta$ CU | SE CU | $\beta$ R1 | SE R1 |
| --- | --- | --- | --- | --- | --- | --- | --- | --- | --- |
| calcarine | LH | 0.137 | 0.109 | 0.173 | 0.132 | -0.197 | 0.206 | -0.122 | 0.076 |
| pos | LH | -0.034 | 0.149 | 0.045 | 0.108 | -0.005 | 0.138 | 0.180 | 0.088 |
| insula | LH | 0.119 | 0.230 | 0.102 | 0.037 | -0.038 | 0.093 | -0.010 | 0.055 |
| central | LH | 0.838 | 0.155 | 0.180 | 0.057 | 0.010 | 0.114 | 0.037 | 0.052 |
| cos | LH | 0.121 | 0.108 | -0.050 | 0.124 | -0.219 | 0.114 | 0.337 | 0.104 |
| sts | LH | 0.762 | 0.140 | 0.051 | 0.074 | -0.260 | 0.112 | 0.022 | 0.081 |
| sfs | LH | 0.160 | 0.054 | 0.226 | 0.089 | -0.275 | 0.119 | 0.110 | 0.060 |
| ips | LH | 0.009 | 0.124 | 0.186 | 0.133 | -0.129 | 0.193 | 0.225 | 0.111 |
| loc | LH | 0.369 | 0.193 | 0.147 | 0.155 | -0.173 | 0.118 | 0.191 | 0.069 |
| ifs | LH | 0.490 | 0.122 | 0.105 | 0.125 | 0.005 | 0.164 | 0.266 | 0.080 |
| ots | LH | 0.103 | 0.078 | -0.216 | 0.095 | -0.416 | 0.108 | 0.349 | 0.069 |
| its | LH | 0.416 | 0.077 | -0.124 | 0.079 | -0.248 | 0.098 | 0.131 | 0.039 |
| calcarine | RH | 0.0329 | 0.0740 | 0.2308 | 0.0894 | 0.1732 | 0.1007 | 0.0684 | 0.0703 |
| pos | RH | 0.1628 | 0.0970 | -0.0943 | 0.0600 | -0.0843 | 0.0824 | 0.1212 | 0.0824 |
| insula | RH | 0.3814 | 0.1572 | 0.0175 | 0.0296 | -0.0801 | 0.0381 | -0.0259 | 0.0501 |
| central | RH | 0.4811 | 0.1173 | 0.1690 | 0.0580 | -0.1969 | 0.0564 | 0.0028 | 0.0510 |
| cos | RH | 0.1540 | 0.1226 | -0.1596 | 0.0951 | -0.1541 | 0.0836 | 0.5992 | 0.1195 |
| sts | RH | 0.6261 | 0.0955 | 0.0415 | 0.0578 | -0.3519 | 0.0481 | -0.0320 | 0.0725 |
| sfs | RH | 0.3304 | 0.0774 | 0.1318 | 0.1070 | -0.0957 | 0.0685 | 0.2557 | 0.0883 |
| ips | RH | 0.0024 | 0.1097 | 0.3263 | 0.1160 | -0.1106 | 0.0715 | 0.2047 | 0.1095 |
| loc | RH | 0.3509 | 0.1112 | -0.0078 | 0.0830 | -0.1995 | 0.1012 | 0.2616 | 0.0728 |
| ifs | RH | 0.3508 | 0.0749 | -0.0223 | 0.0648 | -0.1926 | 0.0629 | 0.2465 | 0.0579 |
| ots | RH | 0.0596 | 0.0807 | -0.1905 | 0.0695 | -0.1437 | 0.0881 | 0.4625 | 0.0668 |
| its | RH | 0.1320 | 0.0510 | 0.0712 | 0.0557 | -0.1603 | 0.0621 | 0.2547 | 0.0553 |

**Supplementary Table 10.** Statistical significance and estimates of the linear mixed model (LMMs): *SD ∼ 1 + (CT + CU + R1) x sulcus + (1|Infant)* across sulci, along the length of each sulcus. Statistics: *DF*: degrees of freedom; *tstats*: t-statistics; *CI* = confidence intervals. *LH/RH:* left/right hemisphere. * = represents an extremely significant and low p-value, which MATLAB reports as *pValue*=0.

| Effects | Hemisphere | $\beta$ Estimate | SE | DF | tstats | pValue | CI Lower | CI Upper |
| --- | --- | --- | --- | --- | --- | --- | --- | --- |
| CT | LH | -0.51 | 0.016 | 45486 | -32.69 | 1.09E-231 | -0.54 | -0.48 |
| CU | LH | 1.01 | 0.011 | 45486 | 93.08 | 0* | 0.99 | 1.03 |
| R1 | LH | 0.09 | 0.016 | 45486 | 5.71 | 1.13E-08 | 0.06 | 0.12 |
| Sulcus | LH | -0.01 | 0.001 | 45486 | -26.99 | 3.37E-159 | -0.01 | -0.01 |
| CT x Sulcus | LH | 0.01 | 0.002 | 45486 | 4.63 | 3.64E-06 | 0.01 | 0.01 |
| CU x Sulcus | LH | -0.05 | 0.002 | 45486 | -30.08 | 7.34E-197 | -0.05 | -0.04 |
| R1 x Sulcus | LH | 0.05 | 0.002 | 45486 | 24.14 | 6.66E-128 | 0.05 | 0.06 |
| CT | RH | -0.16 | 0.014 | 46671 | -11.10 | 1.40E-28 | -0.19 | -0.13 |
| CU | RH | 1.01 | 0.010 | 46671 | 102.75 | 0* | 0.99 | 1.02 |
| R1 | RH | -0.06 | 0.015 | 46671 | -3.93 | 8.61E-05 | -0.09 | -0.03 |
| Sulcus | RH | 0.00 | 0.001 | 46671 | 3.93 | 8.34E-05 | 1.26E-03 | 3.75E-03 |
| CT x Sulcus | RH | -0.03 | 0.002 | 46671 | -13.91 | 6.84E-44 | -0.03 | -0.02 |
| CU x Sulcus | RH | -0.04 | 0.001 | 46671 | -26.57 | 2.15E-154 | -0.04 | -0.03 |
| R1 x Sulcus | RH | 0.05 | 0.002 | 46671 | 25.51 | 1.51E-142 | 0.04 | 0.05 |

**Supplementary Table 11.** Normalized beta coefficients *(β)* and standard errors *(SE)* for cortical thickness (CT), curvature (CU), and cortical microstructure (R_1_) per sulcus in the left/right hemispheres (LH/RH). The beta coefficients represent the relationship between sulcal depth (SD) and each parameter, along the entire length of each sulcus, as described by the model: *SD = β_1_CT + β_2_ CU + β_3_R1 + (1|infant)*.

| Sulcus | Hemi | $\beta$ CT | SE CT | $\beta$ CU | SE CU | $\beta$ R<sub>1</sub> | SE R<sub>1</sub> |
| --- | --- | --- | --- | --- | --- | --- | --- |
| calcarine | LH | -0.10 | 0.03 | 1.07 | 0.02 | -0.05 | 0.03 |
| pos | LH | -0.46 | 0.03 | 0.51 | 0.02 | -0.08 | 0.03 |
| insula | LH | -0.42 | 0.03 | 0.76 | 0.01 | 0.74 | 0.03 |
| central | LH | -0.24 | 0.03 | 0.56 | 0.02 | 0.22 | 0.02 |
| cos | LH | -0.13 | 0.02 | 0.52 | 0.01 | 0.32 | 0.02 |
| sts | LH | -0.57 | 0.03 | 0.64 | 0.01 | 0.67 | 0.03 |
| sfs | LH | -0.59 | 0.03 | 0.65 | 0.01 | 0.73 | 0.03 |
| ips | LH | -0.30 | 0.04 | 0.71 | 0.02 | 0.40 | 0.04 |
| loc | LH | -0.39 | 0.04 | 0.66 | 0.02 | 0.20 | 0.04 |
| ifs | LH | -0.20 | 0.03 | 0.52 | 0.01 | 0.28 | 0.03 |
| ots | LH | -0.25 | 0.03 | 0.57 | 0.02 | 0.84 | 0.04 |
| its | LH | -0.29 | 0.01 | 0.55 | 0.01 | 0.37 | 0.01 |
| calcarine | RH | -0.14 | 0.04 | 1.27 | 0.02 | -0.05 | 0.03 |
| pos | RH | 0.04 | 0.03 | 0.44 | 0.02 | -0.16 | 0.03 |
| insula | RH | -0.55 | 0.03 | 1.04 | 0.01 | 0.28 | 0.04 |
| central | RH | -0.25 | 0.03 | 0.56 | 0.02 | 0.26 | 0.03 |
| cos | RH | -0.36 | 0.02 | 0.68 | 0.01 | 0.76 | 0.02 |
| sts | RH | -0.42 | 0.03 | 0.87 | 0.01 | 0.23 | 0.02 |
| sfs | RH | -0.70 | 0.03 | 0.63 | 0.02 | 0.83 | 0.03 |
| ips | RH | -0.32 | 0.03 | 0.59 | 0.02 | 0.29 | 0.03 |
| loc | RH | -0.54 | 0.04 | 0.74 | 0.02 | 0.22 | 0.04 |
| ifs | RH | -0.45 | 0.03 | 0.46 | 0.02 | 0.73 | 0.04 |
| ots | RH | -0.09 | 0.02 | 0.65 | 0.01 | 0.38 | 0.05 |
| its | RH | -0.31 | 0.01 | 0.63 | 0.01 | 0.22 | 0.01 |

## References

1. Chi, J. G., Dooling, E. C. & Gilles, F. H. Gyral development of the human brain. Ann. Neurol. 1, 86–93 (1977).

2. Garel, C. et al. Fetal cerebral cortex: normal gestational landmarks identified using prenatal MR imaging. AJNR Am. J. Neuroradiol. 22, 184–189 (2001).

3. de Vareilles, H., Rivière, D., Mangin, J. F. & Dubois, J. Development of cortical folds in the human brain: An attempt to review biological hypotheses, early neuroimaging investigations and functional correlates. Dev. Cogn. Neurosci. 61, 101249 (2023).

4. Garcia, K. E., Kroenke, C. D. & Bayly, P. V. Mechanics of cortical folding: stress, growth and stability. Philos. Trans. R. Soc. Lond. B Biol. Sci. 373, (2018).

5. Fernández, V., Llinares-Benadero, C. & Borrell, V. Cerebral cortex expansion and folding: what have we learned? EMBO J. 35, 1021–1044 (2016).

6. Van Essen, D. C. A 2020 view of tension-based cortical morphogenesis. Proc. Natl. Acad. Sci. U. S. A. 117, 32868–32879 (2020).

7. Zilles, K., Palomero-Gallagher, N. & Amunts, K. Development of cortical folding during evolution and ontogeny. Trends Neurosci. 36, 275–284 (2013).

8. Rakic, P. A small step for the cell, a giant leap for mankind: a hypothesis of neocortical expansion during evolution. Trends in Neurosciences 18, 383–388 (1995).

9. Hansen, P. E., Ballesteros, M. C., Soila, K., Garcia, L. & Howard, J. M. MR imaging of the developing human brain. Part 1. Prenatal development. Radiographics 13, 21–36 (1993).

10. Meng, Y., Li, G., Lin, W., Gilmore, J. H. & Shen, D. Spatial distribution and longitudinal development of deep cortical sulcal landmarks in infants. Neuroimage 100, 206–218 (2014).

11. Meng, Y. et al. Discovering cortical sulcal folding patterns in neonates using large-scale dataset. Hum. Brain Mapp. 39, 3625–3635 (2018).

12. Welker, W. Why does cerebral cortex fissure and fold? in Cerebral Cortex 3–136 (Springer US, Boston, MA, 1990).

13. Leroy, F. et al. New human-specific brain landmark: the depth asymmetry of superior temporal sulcus. Proc. Natl. Acad. Sci. U. S. A. 112, 1208–1213 (2015).

14. Natu, V. S. et al. Sulcal Depth in the Medial Ventral Temporal Cortex Predicts the Location of a Place-Selective Region in Macaques, Children, and Adults. Cereb. Cortex 31, 48– 61 (2020).

15. Yao, J. K., Voorhies, W. I., Miller, J. A., Bunge, S. A. & Weiner, K. S. Sulcal depth in prefrontal cortex: a novel predictor of working memory performance. Cereb. Cortex 33, 1799– 1813 (2023).

16. Yun, H. J. et al. Regional Alterations in Cortical Sulcal Depth in Living Fetuses with Down Syndrome. Cereb. Cortex 31, 757–767 (2021).

17. Brun, L. et al. Localized Misfolding Within Broca’s Area as a Distinctive Feature of Autistic Disorder. Biol Psychiatry Cogn Neurosci Neuroimaging 1, 160–168 (2016).

18. Shin, S.-J., Kim, A., Han, K.-M., Tae, W.-S. & Ham, B.-J. Reduced Sulcal Depth in Central Sulcus of Major Depressive Disorder. Exp. Neurobiol. 31, 353–360 (2022).

19. Li, G. et al. Mapping longitudinal development of local cortical gyrification in infants from birth to 2 years of age. J. Neurosci. 34, 4228–4238 (2014).

20. Im, K. & Grant, P. E. Sulcal pits and patterns in developing human brains. Neuroimage 185, 881–890 (2019).

21. Holland, M. A. et al. Folding drives cortical thickness variations. Eur. Phys. J. Spec. Top. 229, 2757–2778 (2020).

22. Demirci, N. & Holland, M. A. Cortical thickness systematically varies with curvature and depth in healthy human brains. Hum. Brain Mapp. 43, 2064–2084 (2022).

23. Hill, J. et al. A surface-based analysis of hemispheric asymmetries and folding of cerebral cortex in term-born human infants. J. Neurosci. 30, 2268–2276 (2010).

24. Li, G. et al. Mapping longitudinal hemispheric structural asymmetries of the human cerebral cortex from birth to 2 years of age. Cereb. Cortex 24, 1289–1300 (2014).

25. Shimony, J. S. et al. Comparison of cortical folding measures for evaluation of developing human brain. Neuroimage 125, 780–790 (2016).

26. Duan, D. et al. Exploring folding patterns of infant cerebral cortex based on multi-view curvature features: Methods and applications. Neuroimage 185, 575–592 (2019).

27. Ahmad, S. et al. Multifaceted atlases of the human brain in its infancy. Nat. Methods 20, 55–64 (2023).

28. Bethlehem, R. A. I. et al. Brain charts for the human lifespan. Nature 604, 525–533 (2022).

29. Rakic, P., Bourgeois, J. P. & Goldman-Rakic, P. S. Synaptic development of the cerebral cortex: implications for learning, memory, and mental illness. Prog. Brain Res. 102, 227–243 (1994).

30. Huttenlocher, P. R. & Dabholkar, A. S. Regional differences in synaptogenesis in human cerebral cortex. J. Comp. Neurol. 387, 167–178 (1997).

31. Elston, G. N., Oga, T. & Fujita, I. Spinogenesis and pruning scales across functional hierarchies. J. Neurosci. 29, 3271–3275 (2009).

32. Elston, G. N. & Fujita, I. Pyramidal cell development: postnatal spinogenesis, dendritic growth, axon growth, and electrophysiology. Front. Neuroanat. 8, 78 (2014).

33. Miller, D. J. et al. Prolonged myelination in human neocortical evolution. Proc. Natl. Acad. Sci. U. S. A. 109, 16480–16485 (2012).

34. Mezer, A. et al. Quantifying the local tissue volume and composition in individual brains with magnetic resonance imaging. Nat. Med. 19, 1667–1672 (2013).

35. Edwards, L. J., Kirilina, E., Mohammadi, S. & Weiskopf, N. Microstructural imaging of human neocortex in vivo. Neuroimage 182, 184–206 (2018).

36. Weiskopf, N., Edwards, L. J., Helms, G., Mohammadi, S. & Kirilina, E. Quantitative magnetic resonance imaging of brain anatomy and in vivo histology. Nature Reviews Physics 3, 570–588 (2021).

37. Möller, H. E. et al. Iron, Myelin, and the Brain: Neuroimaging Meets Neurobiology. Trends Neurosci. 42, 384–401 (2019).

38. Stüber, C. et al. Myelin and iron concentration in the human brain: a quantitative study of MRI contrast. Neuroimage 93 Pt 1, 95–106 (2014).

39. Natu, V. S. et al. Apparent thinning of human visual cortex during childhood is associated with myelination. Proc. Natl. Acad. Sci. U. S. A. 116, 20750–20759 (2019).

40. Natu, V. S. et al. Infants’ cortex undergoes microstructural growth coupled with myelination during development. Commun Biol 4, 1191 (2021).

41. Richman, D. P., Stewart, R. M., Hutchinson, J. & Caviness, V. S. Mechanical Model of Brain Convolutional Development. Science 189, 18–21 (1975).

42. Toro, R. & Burnod, Y. A morphogenetic model for the development of cortical convolutions. Cereb. Cortex 15, 1900–1913 (2005).

43. Kriegstein, A., Noctor, S. & Martínez-Cerdeño, V. Patterns of neural stem and progenitor cell division may underlie evolutionary cortical expansion. Nat. Rev. Neurosci. 7, 883–890 (2006).

44. Reillo, I., de Juan Romero, C., García-Cabezas, M. Á. & Borrell, V. A role for intermediate radial glia in the tangential expansion of the mammalian cerebral cortex. Cereb. Cortex 21, 1674–1694 (2011).

45. Zöllei, L., Iglesias, J. E., Ou, Y., Grant, P. E. & Fischl, B. Infant FreeSurfer: An automated segmentation and surface extraction pipeline for T1-weighted neuroimaging data of infants 0-2 years. Neuroimage 218, 116946 (2020).

46. Fischl, B., Sereno, M. I., Tootell, R. B. & Dale, A. M. High-resolution intersubject averaging and a coordinate system for the cortical surface. Hum. Brain Mapp. 8, 272–284 (1999).

47. Yousry, T. A. et al. Localization of the motor hand area to a knob on the precentral gyrus. A new landmark. Brain 120 (Pt 1), 141–157 (1997).

48. Coulon, O. et al. Two new stable anatomical landmarks on the Central Sulcus: definition, automatic detection, and their relationship with primary motor functions of the hand. Annu. Int. Conf. IEEE Eng. Med. Biol. Soc. 2011, 7795–7798 (2011).

49. Bodin, C., Takerkart, S., Belin, P. & Coulon, O. Anatomo-functional correspondence in the superior temporal sulcus. Brain Struct. Funct. 223, 221–232 (2018).

50. Eichert, N., Watkins, K. E., Mars, R. B. & Petrides, M. Morphological and functional variability in central and subcentral motor cortex of the human brain. Brain Struct. Funct. 226, 263–279 (2021).

51. Bartha-Doering, L. et al. Fetal temporal sulcus depth asymmetry has prognostic value for language development. *Commun*. Biol. 6, 109 (2023).

52. Vandekar, S. N. et al. Topologically dissociable patterns of development of the human cerebral cortex. J. Neurosci. 35, 599–609 (2015).

53. Dubois, J. et al. Mapping the early cortical folding process in the preterm newborn brain. Cereb. Cortex 18, 1444–1454 (2008).

54. Yun, H. J. et al. Temporal patterns of emergence and spatial distribution of sulcal pits during fetal life. Cereb. Cortex 30, 4257–4268 (2020).

55. Rakic, P. Neuroscience. Genetic control of cortical convolutions. Science vol. 303 1983–1984 (2004).

56. Tallinen, T., Chung, J. Y., Biggins, J. S. & Mahadevan, L. Gyrification from constrained cortical expansion. Proc. Natl. Acad. Sci. U. S. A. 111, 12667–12672 (2014).

57. Fitzgibbon, S. P. et al. The developing Human Connectome Project (dHCP) automated resting-state functional processing framework for newborn infants. Neuroimage 223, 117303 (2020).

58. Howell, B. R. et al. The UNC/UMN Baby Connectome Project (BCP): An overview of the study design and protocol development. Neuroimage 185, 891–905 (2019).

59. Casey, B. J. et al. The Adolescent Brain Cognitive Development (ABCD) study: Imaging acquisition across 21 sites. Dev. Cogn. Neurosci. 32, 43–54 (2018).

60. Dubois, J. et al. Primary cortical folding in the human newborn: an early marker of later functional development. Brain 131, 2028–2041 (2008).

61. Wandell, B. A., Dumoulin, S. O. & Brewer, A. A. Visual field maps in human cortex. Neuron 56, 366–383 (2007).

62. Katz, L. C. & Shatz, C. J. Synaptic activity and the construction of cortical circuits. Science 274, 1133–1138 (1996).

63. Long, B., et al. The BabyView dataset: High-resolution egocentric videos of infants’ and young children’s everyday experiences. arXiv [cs.CV] (2024).

64. Zhuang, C. et al. Unsupervised neural network models of the ventral visual stream. Proc. Natl. Acad. Sci. U. S. A. 118, e2014196118 (2021).

65. Turk-Browne, N. B. & Aslin, R. N. Infant neuroscience: how to measure brain activity in the youngest minds. Trends Neurosci. 47, 338–354 (2024).

66. Hasson, U., Nastase, S. A. & Goldstein, A. Direct fit to nature: An evolutionary perspective on biological and artificial neural networks. Neuron 105, 416–434 (2020).

67. Smith, L. B., Jayaraman, S., Clerkin, E. & Yu, C. The developing infant creates a curriculum for statistical learning. Trends Cogn. Sci. 22, 325–336 (2018).

68. O’Doherty, C. Deep neural network models of infant visual cortex. https://2024.ccneuro.org/pdf/380_Paper_authored_CCN_2024.pdf.

69. Ellis, C. T. et al. Retinotopic organization of visual cortex in human infants. Neuron 109, 2616–2626.e6 (2021).

70. Kosakowski, H. et al. Selective responses to faces, scenes, and bodies in the ventral visual pathway of infants. PsyArXiv (2021) doi:10.31234/osf.io/7hqcu.

71. Ramsaran, A. I. & Frankland, P. W. Babies form fleeting memories. Science (New York, N.Y.) vol. 387 1253–1254 (2025).

72. Deen, B. et al. Organization of high-level visual cortex in human infants. Nat. Commun. 8, 13995 (2017).

73. Nordahl, C. W. et al. Cortical folding abnormalities in autism revealed by surface-based morphometry. J. Neurosci. 27, 11725–11735 (2007).

74. Csernansky, J. G. et al. Symmetric abnormalities in sulcal patterning in schizophrenia. Neuroimage 43, 440–446 (2008).

75. Ghotra, A. et al. A size-adaptive 32-channel array coil for awake infant neuroimaging at 3 Tesla MRI. Magn. Reson. Med. 86, 1773–1785 (2021).

76. Jenkinson, M., Beckmann, C. F., Behrens, T. E. J., Woolrich, M. W. & Smith, S. M. FSL. Neuroimage 62, 782–790 (2012).

77. Wang, L. et al. iBEAT V2.0: a multisite-applicable, deep learning-based pipeline for infant cerebral cortical surface reconstruction. Nat. Protoc. 18, 1488–1509 (2023).

78. Bosker, R. & Snijders, T. Multilevel Analysis : Introduction Basic Advanced Multilevel Modeling. 1–368 (2011).

